# Bcl-xL restricts transcriptional, morphological and functional decompensation of β-cell mitochondria under chronic glucose excess

**DOI:** 10.1101/2021.10.25.465491

**Authors:** Daniel J. Pasula, Rocky Shi, Ben Vanderkruk, Alexis Z.L. Shih, Yuanjie Zou, Ahsen Chaudhry, Brad G. Hoffman, Dan S. Luciani

## Abstract

In the progression of diabetes, pancreatic islet β-cells respond to increased metabolic demand with functional compensation, followed by pathogenic decompensation of mitochondria-dependent insulin secretion. It is not clear what mechanisms drive, or control, mitochondrial decompensation. Here, we report that anti-apoptotic Bcl-x_L_ maintains mitochondrial integrity in β-cells under non-apoptotic levels of glucose stress. Prolonged glucose excess causes transcriptional reprogramming of glycolysis and β-cell identity genes, while sensitizing glucose-stimulated Ca^2+^ signaling and insulin secretion. Deletion of Bcl-x_L_ amplifies this insulin hypersecretion and increases mitochondrial fusion, mitochondrial volume, and oxygen consumption, whereas ATP-coupled respiration and mitochondrial hyperpolarization become impaired. Of note, Bcl-x_L_-deficient β-cells have impaired Pgc-1α expression, and develop specific defects in the expression of *Tfam,* mitochondrial ribosomal genes, and OXPHOS components under glucose stress. Bcl-x_L_ limits high glucose-induced mitochondrial ROS (mitoROS) levels and pharmacological normalization of mitoROS in Bcl-x_L_ KO cells rescues glucose-induced defects in mitochondrial gene expression and changes to β-cell identity. Our data identify mitoROS as a primary retrograde driver of transcriptional re-wiring in β-cells exposed to excess glucose, and reveal Bcl-x_L_ as an important safeguard against transcriptional and functional decompensation of β-cell mitochondria. Bcl-x_L_ and mitoROS may thus be viable targets to prevent early β-cell dysfunction and the progression of diabetes.

## INTRODUCTION

Nutrient-induced β-cell insulin secretion is essential to maintain normoglycemia, and β-cell failure is a defining event in the development of type 2 diabetes (T2D). In response to worsening insulin resistance and nutrient excess β-cells respond by increasing their total insulin output, which may compensate sufficiently to maintain euglycemia and prevent the progression toward prediabetes and diabetes. Evidence suggests that β-cell compensation involves both an initial augmentation of cellular function^1,2^, and a slower expansion of total β-cell mass^2^. However, in individuals that progress to develop diabetes, the compensatory response is transient due to a progressive development of β-cell dysfunction, and ultimately also a loss of β-cells by dedifferentiation, trans-differentiation and apoptosis.

β-Cell mitochondrial metabolism generates ATP and other metabolic coupling factors that are essential for glucose-stimulated insulin secretion^3^. Severe disruptions to both mitochondrial function and structure are seen in β-cells from mice and humans with T2D, indicating that mitochondrial dysfunction plays a role in the failure of nutrient-induced insulin secretion^4–6^. Recent findings have shown that relatively modest increases in blood glucose are associated with changes in the expression of genes related to β-cell identity and metabolism^7^, and that the progression of diabetes and hyperglycemia is associated with increasing disruption of transcripts and proteins involved in oxidative phosphorylation and other aspects of mitochondrial physiology^8^. Comparatively little is known about the mechanisms that mediate the earlier stages of functional adaptation, but augmented mitochondrial metabolism likely plays a central role^9^. Therefore, glucose-induced changes to β-cell mitochondrial function appear central to the development of T2D. Despite these recent insights, it remains unclear what mechanisms drive the progression from mitochondrial adaptation to failure under conditions of chronic glucose excess.

Apoptosis-regulating proteins in the Bcl-2 family have emerged as important regulators of physiological processes in non-transformed cells, including notable roles of both pro- and anti-apoptotic proteins in the regulation of cellular metabolism^10^. We, and others, have linked Bcl-2 family proteins to glucose-stimulated mitochondrial function and insulin secretion in pancreatic β-cells^11–14^. Our previous work demonstrated that anti-apoptotic Bcl-x_L_ dampens the responsiveness of β-cell mitochondria to glucose stimulation^12^, and we hypothesized that these non-canonical metabolic functions could play an important role in maintaining mitochondrial integrity in β-cells during increases in metabolic demand.

In this paper, we tested this possibility by examining the relationship between β-cell gene expression, function, and mitochondrial homeostasis in Bcl-x_L_ wild-type and knockout β-cells exposed to chronic high levels of glucose. Our results reveal a previously unrecognized role of Bcl-x_L_ in limiting the progression of mitochondrial decompensation in β-cells under non-apoptotic levels of metabolic stress.

## METHODS

### Reagents

Tamoxifen (#T5648), Collagenase type XI (#C7657), Tetramethylrhodamine Ethyl Ester (TMRE, #87917), D-glucose (#G7528), Diazoxide (DM, #D9035), MitoTEMPO (#SML0737), FCCP (#C2920), Oligomycin (#O4876), Antimycin-A (#A8674) and Rotenone (#R8875) were purchased from Sigma-Aldrich (St. Louis, MO). Fura-2 AM (#F1221), MitoTracker Green FM (MTG, #M7514), Hoechst 33342 (#H3570), MitoSOX (#M36008), RPMI 1640 (#11879), Dulbecco’s Modified Eagle’s Medium (DMEM, #11995), Fetal Bovine Serum (FBS, #10438), Trypsin-EDTA (#25300), Penicillin-Streptomycin 10,000 U/ml (#15140), and HBSS (#14185) were from Life Technologies/Thermo Fisher Scientific (Carlsbad, CA). Dimethyl sulfoxide (DMSO, #BP231) was purchased from Fisher Scientific (Waltham, MA) and Minimum Essential Media (MEM, #15-015-CV) was from Corning (Corning, NY). Seahorse XF^e^24 Islet Capture FluxPak (103518-100) were purchased from Agilent Technologies Canada (Mississauga, ON).

### Animals

Mice with tamoxifen (TM)-inducible β-cell-selective deletion of Bcl-x_L_ were established and bred as previously described^12^. Pdx1-CreER^TM^:Bclx^fl/fl^ mice were injected 4 consecutive days with 3 mg/40g TM injection to induce gene deletion and generate Bcl-x_L_ knockout (BclxβKO) mice. TM-injected littermate Bclx^fl/fl^ (BclxβWT) mice were used as controls to account for possible metabolic effects of tamoxifen^15,16^. To avoid potential long-term compensation for Bcl-x_L_ deletion, islets were isolated 5-7 days after the last TM injection. Experiments were done using 14-16 week old male mice. All animal procedures were done according to national and international guidelines and approved by the University of British Columbia Animal Care Committee.

### Islet Isolation and Cell Culture

Pancreatic islets were isolated by our previously described methods of collagenase digestion and filtration-based purification^11^. The isolated islets were hand-picked and allowed to rest overnight before further analysis or experimental cultures. Aliquots of islets from all experimental mice were collected for qPCR confirmation of tamoxifen-induced gene deletion. For single cell microscopy the islets were dispersed and seeded on 25 mm glass coverslips^17^. The effects of nutrient excess were examined by culturing intact islets or dispersed islet-cells for 6 days in RPMI completed with 10% FBS and 2% Penicillin-Streptomycin and containing either 11 mM glucose (normal glucose; NG) or 25 mM glucose (high glucose; HG).

### Respiratory Control Analysis

Islet respiration was quantified using the Seahorse XF^e^24 Analyzer (Agilent Technologies). Intact islets were washed with PBS twice and loaded into XF^e^24 islet capture microplates in Seahorse XF RPMI Medium pH 7.4 (#103576) supplemented with 3 mM glucose, 2 mM Glutamine and 2 mM Sodium Pyruvate and incubated at 37°C in a non-CO_2_ incubator for 60 min before loading into the XF^e^24 Analyzer. After the basal oxygen consumption rate (OCR) measurement reached a steady state, the wells were injected with either – 1) increasing concentrations of glucose followed by 1 µM oligomycin; or 2) exposed to a mitochondrial stress test, consisting of the sequential injection of 15 mM glucose, 1 µM oligomycin, 2 µM FCCP and finally a combination of 1 µM rotenone and 1 µM antimycin A. The amount of ATP-coupled respiration, ATP coupling efficiency, proton leak and spare respiratory capacity were calculated as described in^18^. OCR values were normalized to total DNA per well quantified on Qubit^TM^ Flurometer (Q32857 Invitrogen) using Qubit^TM^ dsDNA HS Assay Kit (Q32854).

### Ca^2+^ Imaging

Intact islets were pre-cultured 2-6 days on glass coverslips in RPMI containing glucose concentrations as indicated. Cytosolic Ca^2+^ signaling was then measured using Fura-2AM fluorescence microscopy, essentially as previously^11^. Briefly, islets were loaded with 5 µM Fura-2 for 30 min in RPMI containing 3 mM glucose and transferred to a 2 ml imaging chamber for continuous perifusion at 2.5 ml/min with Ringer’s solution containing 5.5 mM KCl, 2 mM CaCl_2_, 1 mM MgCl_2_, 20 mM HEPES, and 144 mM NaCl and varying glucose concentrations. Before image acquisition, the perifused islets were equilibrated for 30 min with 3 mM glucose Ringer’s and Ca^2+^ changes then recorded in response to glucose and other stimuli, as indicated. Cytosolic Ca^2+^ levels are expressed as the Fura-2 fluorescence intensity ratio (F_340_/F_380_).

### Confocal Imaging and Analysis of Mitochondria

Live islet-cells were imaged in a Tokai Hit INUBTFP-WSKM stage-top incubator at 37°C and 5% CO_2_ on a Leica SP8 Laser Scanning Confocal Microscope (Concord, Ontario, Canada). Optimized 2D and 3D confocal image acquisition, as well as quantitative extraction of mitochondrial features, was done according to our recent pipeline for mitochondrial analysis^19^. For automated batch analysis we used the associated Mitochondria Analyzer plugin that we developed for the ImageJ/Fiji and have made available online^20,21^.

### Machine Learning-based Classification of Mitochondrial Morphology and Networking

For machine learning-based classification of 3D mitochondrial features we first generated a training set based on image stacks of MTG-labelled mouse islet-cells and MIN6 cells expressing mitochondria-targeted YFP. From these images, morphological and networking descriptors were quantified for a total of 2190 mitochondria and the data was grouped into 7 clusters by unsupervised K-means cluster analysis in the XLSTAT software (Addinsoft, NY). The clusters were visually examined and merged based on similarity until 4 major groupings remained, which we classified as Punctate, Tubular, Filamentous, and Highly Complex based on their visual characteristics. The principal features of each morphological classification are summarized in Table 1. All 2190 mitochondria were then manually inspected to verify their correct classification and reassignments were made as necessary. The final training set was then produced by re-quantifying the morphological and networking descriptors of all mitochondria within each of the classifications.

**Table 1:** Description of features that characterize mitochondrial morphological classifications.

| Classification | Features |
| --- | --- |
| Punctate | Spheroid, small mitochondria with little to no branch length or complexity. |
| Tubular | Long or elongated mitochondria with variable branch length, but only 1 or few branches. |
| Filamentous | Networked mitochondria with multiple branches. |
| Highly Complex | Extremely connected mitochondria with very high branch descriptor values. Typically $\geq 8\mu\text{m}^3$ volume or containing >10% of cell's total mitochondrial volume. |

After establishing the training set, it was used to classify mitochondria of experimental cells using a Random Forest Classification algorithm in XLSTAT. The mitochondrial profile of a cell was then established as the proportion of each mitochondrial “morpho-types” normalized to the total mitochondrial volume in the cell. A summary of the workflow is shown in Supplemental Fig. 1.

### Mitochondrial ROS Imaging

Dispersed islet cells were pre-cultured for 6 days on glass coverslips in either NG or HG RPMI with or without addition of 500 nM of the mitochondria-targeted superoxide scavenger MitoTEMPO. Mitochondrial ROS was then measured using MitoSOX Red superoxide indicator. Briefly, cells were washed and then stained with 5 µM MitoSOX in phenol red free RPMI protected from light and incubated at 37°C and 5% CO_2_ for 15 minutes before being imaged using a Leica DMI6000 inverted microscope equipped with an HCX Plan FLUOTAR L 20x objective, a DFC365FX digital camera, 546/12x excitation and 605/75m emission filters.

### Islet Insulin Secretion and Insulin Content

Insulin secretion was measured under static incubations from 15 size-matched islets. Following 6-day pre-culture, the islets were collected in 1.5 ml low retention microcentrifuge tubes and equilibrated for 1 hour in Kreb’s Ringer’s buffer (KRB; 129 mM NaCl, 4.8 mM KCl, 1.2 mM MgSO_4_, 1.2 mM KH_2_PO_4_, 2.5 mM CaCl_2_, 5 mM NaHCO_3_, 10 mM HEPES and 0.5% BSA) with 3 mM glucose. The same groups of islets were then transferred between tubes containing 500 µl of KRB with glucose and treatments as indicated, moving sequentially from low to high glucose conditions. After each 60 min incubation step the islets were spun down at 4°C, supernatant was collected for measurement of secreted insulin, and the islets were re-counted before transfer to the next tube. Islet insulin content was extracted by washing twice with chilled PBS and freeze-thawing in 100 μl of RIPA buffer at -20°C before assay. Insulin concentrations were measured using the Ultrasensitive Insulin ELISA kit (Alpco #80-INSMSU-E10) according to manufacturer’s manual. Luminescence was measured at 450 nm on a SpectraMaxL luminometer (Molecular Devices)

### RT-qPCR

For each sample, 40 to 50 islets were preserved in 40 μl RNAlater (Qiagen #76104) at -80°C before RNA isolation. Total RNA was isolated using the RNEasy Mini Kit (Qiagen #74106) and measured by NanoDropTM 2000 (Thermo Fisher, Carlsbad, California). Freshly isolated RNA (100 ng/sample) was reverse transcribed using qScript cDNA synthesis kit (Quanta Bioscience #95047-500) and cDNA was stored at -20°C until use. The qPCR reactions were run using: 4 μl PerfeCTa SYBR green Fastmix (Quanta Biosciences #95072), 5 μM forward and reverse mixed primers, 1 μl cDNA, and 1.5 μl ddH_2_O. Primer efficiencies were validated by serial dilutions and amplicon size confirmed by agarose gel electrophoresis. Validated primer sequences are listed in Table 2. Expression levels were assayed in triplicates on 384-well microplates (Life technologies #4309849) on a ViiA7 Real-Time PCR machine (Applied Biosystems, Foster City, California) and normalized to the *Actb* housekeeping gene.

**Table 2:**
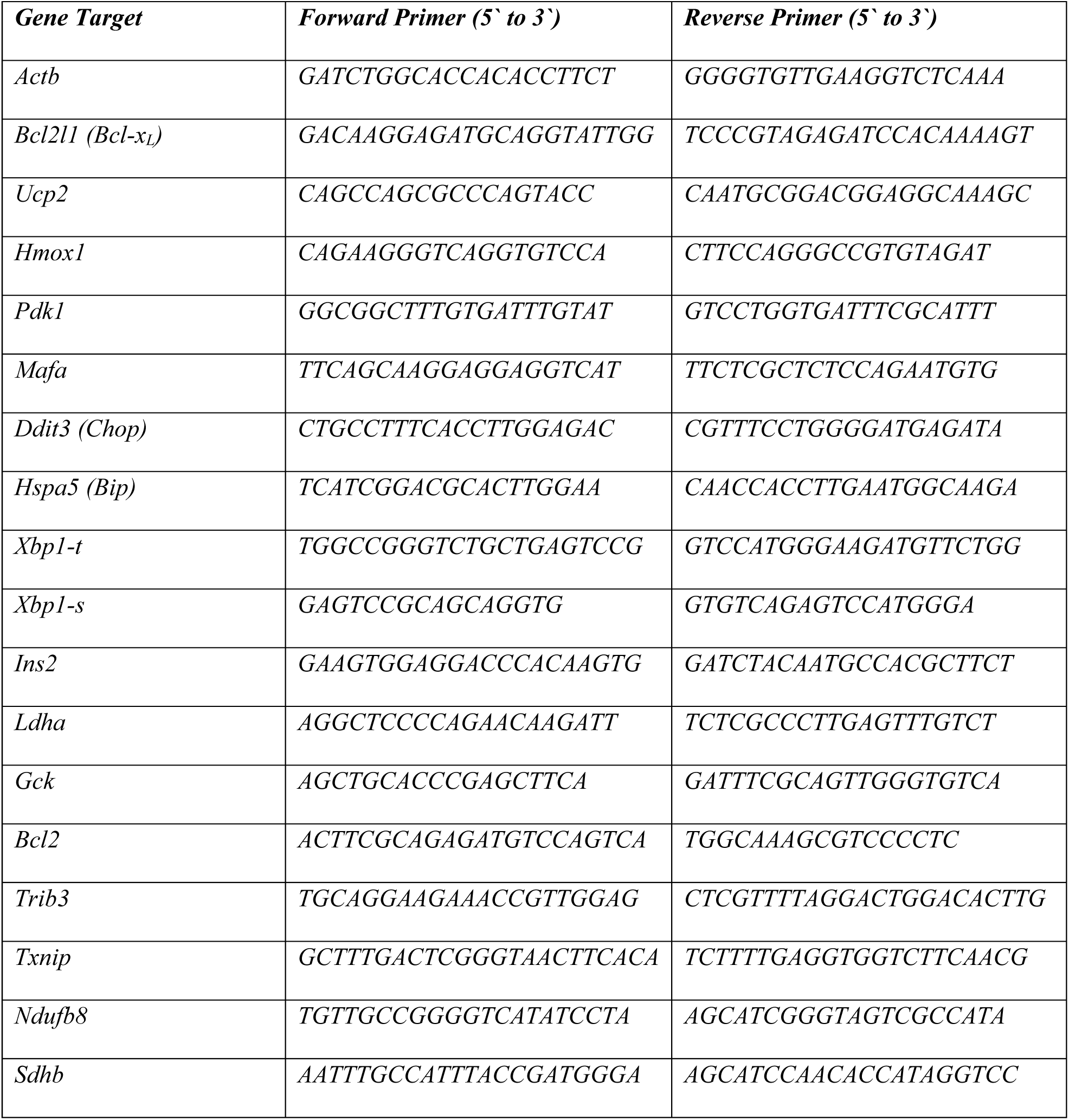
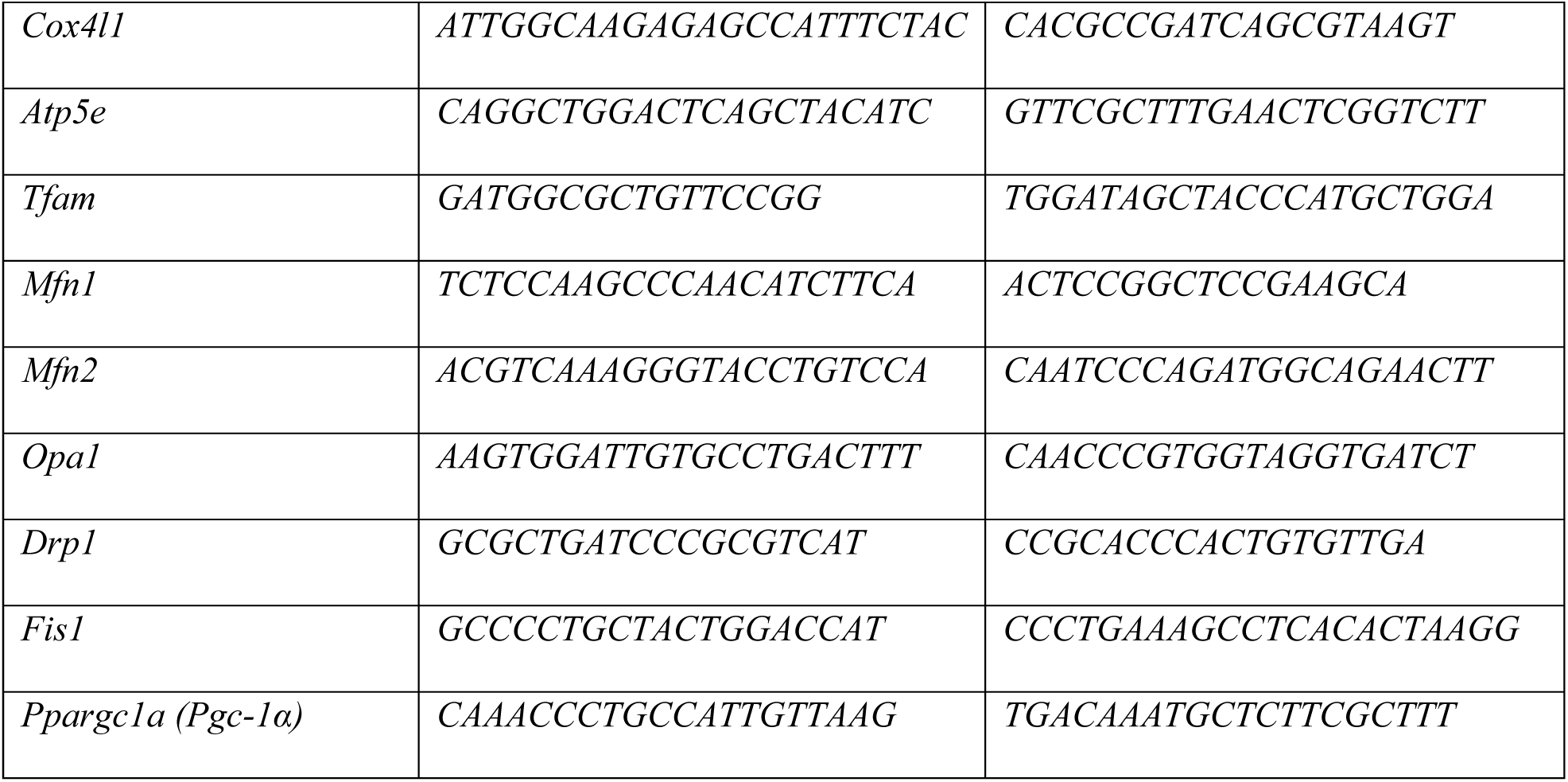
qPCR primers used

### RNA-Sequencing

#### Sample preparation and sequencing

RNA was extracted from whole pancreatic islets using Trizol Reagent (Life Technologies, 15596026) and ethanol precipitation with glycogen as a carrier, followed by DNase-treatment using Turbo DNA Free Kit (Life Technologies, AM1907). Total RNA content was estimated using Nanodrop, and samples were prepared for sequencing from 400 ng of total RNA using NEBNext Ultra II Directional RNA Library Kit for Illumina with poy(A) mRNA enrichment (NEB, E7760, E7490) according to the manufacturer’s protocols. Libraries were quantified using Qubit fluorometer and Agilent Bioanalyzer, pooled, and sequenced on an Illumina NextSeq 500 (paired-end, 38 base-pair reads) to a depth of approximately 30 million reads per sample.

#### Data alignment

Quality of the sequenced RNA-Seq libraries was assessed using FastQC v0.11.2 (Babraham Bioinformatics). Read quality filtering and adapter trimming were carried out using Trimmomatic v0.39 ^22^. The resulting reads were then aligned and quantified at the transcript level using Salmon^23^ against GENCODE M20/GRCm38.p6 with the default parameters, and aggregated to the gene level using tximport^24^.

#### Differential gene expression analysis and quantification

Gene expression values measured as relative counts were generated using DESeq2 v1.24.0 ^25^, and a minimum threshold of 10 gene counts across all samples was imposed. Differentially-expressed genes were identified in pair-wise comparisons of the four biological conditions (WT_NG; WT_HG; KO_NG; KO_HG) using DESeq2, implementing the *apeglm* method^26^ for effect-size estimation. Genes with fold-change ≥ 1.5 and *p*_adj_ ≤ 0.05 were considered significantly down- or up-regulated. Data visualizations were generated using custom R scripts.

#### Gene-ontology analysis

Biological pathway and function analyses were performed using The Database for Annotation, Visualization and Integrated Discovery (DAVID v6.8) with default parameter settings^27,28^.

### Statistical Analysis

All data were represented as mean ± standard error of the mean (SEM). Data were analyzed in GraphPad Prism 9.0 (La Jolla, California) using Student’s t-test, one-way ANOVA, or two-way ANOVA followed by multiple comparison tests, as appropriate. Statistical significance was set at a threshold of *p* < 0.05.

## RESULTS

### Islet transcriptional adaptation during prolonged high glucose exposure

To examine the effects of sustained metabolic demand, we used a controlled *in vitro* model consisting of 6-day islet culture in normal glucose media (11 mM glucose; NG) or high glucose media (25 mM glucose; HG) (Fig. 1a). For a broad overview of high glucose-induced changes to islet gene expression, we performed bulk RNA-Seq analysis of NG- and HG-cultured BclxβWT and BclxβKO islets. High glucose caused major changes to the transcriptional profiles; in BclxβWT islets the expression of 1708 genes (654 down and 1054 up) were altered, and in BclxβKO islets 2182 genes (1013 down and 1169 up) were differentially expressed after HG culture. Of these HG-sensitive transcripts, 725 were altered in both genotypes (Fig. 1b). Focusing first on the common, genotype-independent, response to excess glucose, we performed pathway enrichment analysis, which revealed that the shared transcripts were dominated by changes related to protein and amino acid processing, as well as cellular metabolism (Fig. 1b). Notably, the strongest enrichment was associated with protein processing in the endoplasmic reticulum (ER). Glucose-stimulated insulin biosynthesis increases pressure on the ER protein processing and folding machinery, which can lead to ER stress and apoptosis if the system is overwhelmed. To evaluate islet ER stress status, we compared the shared transcripts to those associated with the gene ontology term “Response to ER stress” (GO:0034976). Of the 725 transcripts, 32 fell in this category and together they indicated a robust up-regulation related to adaptive ER protein homeostasis, including ER-associated degradation (ERAD), protein translation and folding (Fig. 1c). *Itpr1*, which encodes isoform 1 of the IP_3_R ER Ca^2+^ release channel was down-regulated in HG culture. We previously demonstrated that IP_3_R activity can exacerbate β-cell ER stress and apoptosis^29^, suggesting that *Itpr1* down-regulation is a protective ER response. The pro-apoptotic gene *Trib3* was induced in both BclxβWT and BclxβKO islets and HG culture also caused some genotype-specific changes to ER stress genes, but neither genotype showed other defining characteristics of the transition toward ER stress-induced β-cell apoptosis, such as loss of the adaptive unfolded protein response (UPR) or significant induction of *Ddit3* (aka *Chop*)^30^ (Fig. 1e & Supplemental Fig. 2).

**Figure 1.**
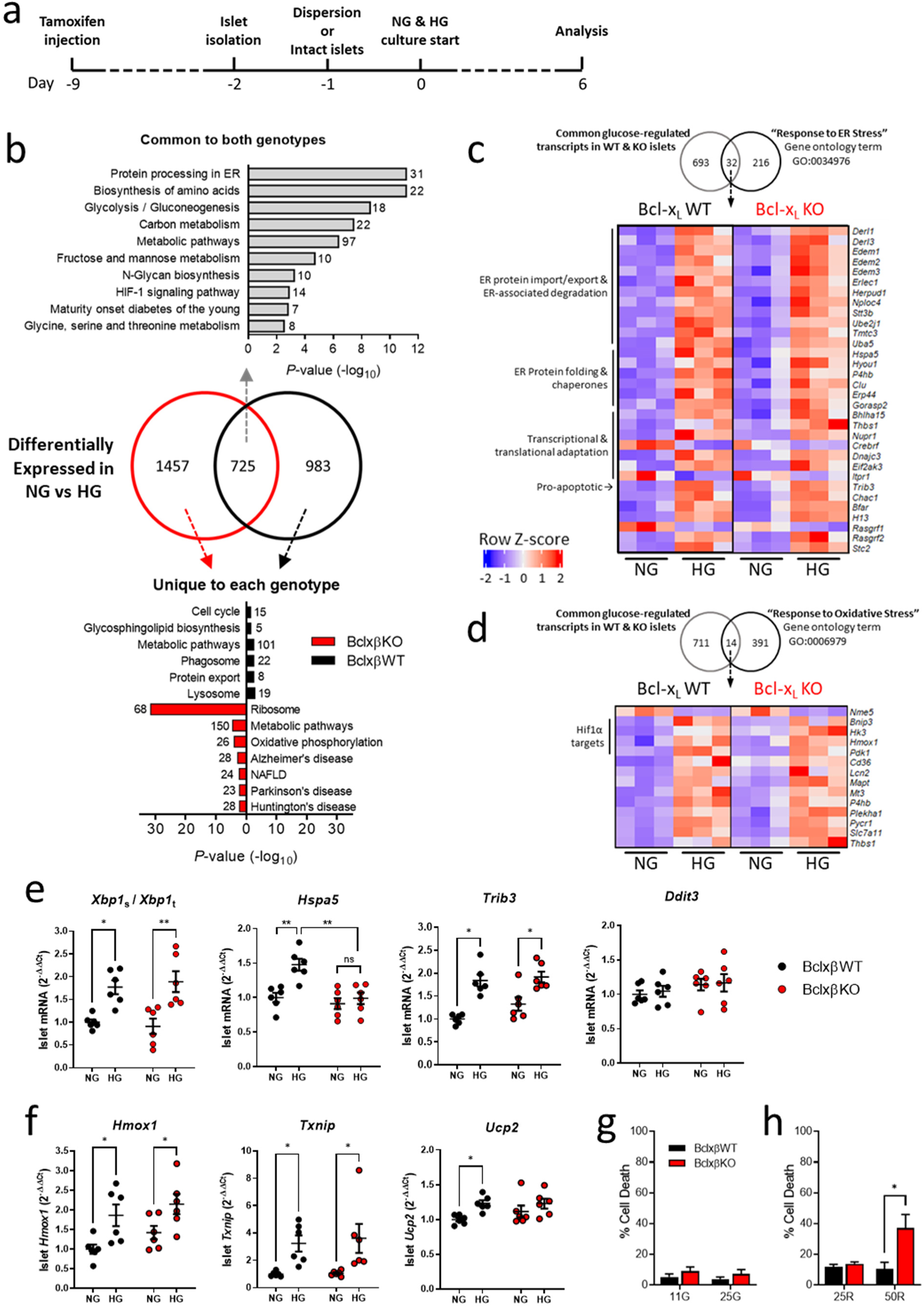
Effects of prolonged high glucose culture on islet transcriptome, stress pathways and survival. a) Schematic illustration of protocol for metabolic challenge experiment. NG = normal glucose (11 mM glucose RPMI media), HG = high glucose (25 mM glucose RPMI media). b) Pathway enrichment analysis of transcripts that were altered in HG-cultured islets compared to NG-cultured islets from BclxβWT and BclxβKO mice. *Top grey*: Pathways associated with transcripts that were differentially expressed in both genotypes. *Bottom:* Pathways associated with transcripts that were uniquely affected in one of the genotypes. Black = BclxβWT, Red = BclxβKO. The number of transcripts in each category is indicated by each bar (n=3 independent islet cDNA library preparations). c) Heat map of genes that were differentially expressed in both BclxβWT and BclxβKO islets after HG culture and are associated with the Gene Ontology term “Cellular response to ER stress” (GO:003497). d) Heat map of genes that were differentially expressed in both BclxβWT and BclxβKO islets after HG culture and are associated with the Gene Ontology term “Oxidative Stress Response” (GO: 0006979). e) Select ER stress-related gene expression changes quantified by qPCR (n=6). f) Select oxidative-stress related gene expression changes quantified by qPCR (n=6). g) Death of Bcl-x_L_ WT and Bclx_L_ KO cells after islet culture in 11 mM glucose (11G) or 25 mM glucose (25G) for 6 days (n=3). No significant differences by two-way ANOVA. h) Death of Bcl-x_L_ WT and Bcl-x_L_ KO cells after culture in 25 mM ribose (25R) or 50 mM ribose (50R) for 4 days (n=5). In all panels, *\*p*<0.05, \*\**p*<0.01 by two-way ANOVA followed by Šidák’s multiple comparison.

β-Cell glucotoxicity is also mediated by oxidative stress, so we also compared common HG-altered transcripts to the gene ontology term “Oxidative Stress Response” (GO: 0006979). In combination with qPCR, this identified some ROS-associated changes to islet mRNA expression; *Hmox1* and *Txnip* were up-regulated but other major redox-regulated transcripts, including *Gpx1, Cat, Sod1, Sod2 and Nfe2l2* were not significantly affected (Fig. 1d,f and not shown). This indicates some activation of oxidative stress, but not sufficiently severe to induce large-scale changes to the expression of antioxidant response genes.

Further suggesting that the high glucose challenge was not overtly toxic, the RNA-Seq analysis did not show a pro-apoptotic shift in Bcl-2 family transcripts^31^. Major family members were unaltered by HG in BclxβWT islets, whereas BclxβKO islets only showed an increase in anti-apoptotic *Bcl2l2* (aka *Bclw; p*_adj_ < 0.01, NG vs HG). Moreover, no significant islet cell death was detected in response to the 6-day high glucose culture, even in the absence of Bcl-x_L_ (Fig. 1g). Using a more toxic challenge with high concentrations of ribose^32–34^, we established that deletion of Bcl-x_L_ did in fact sensitize β-cells to oxidative stress-related death (Fig. 1h), consistent with its canonical function in restricting Bax- and Bak-mediated apoptosis^35^.

Overall, this shows that in our model of prolonged glucose excess, the common transcriptional changes in BclxβWT and BclxβKO islets include signs of beginning oxidative stress, but, as a whole, are non-apoptotic and dominated by robust activation of the adaptive UPR.

### Effects of high glucose culture on the expression of glycolytic and β-cell identity genes

In addition to ER protein processing, both BclxβWT and BclxβKO islets responded to HG with altered expression of metabolic genes (Fig. 1). The common changes included up-regulation of a number of transcripts in glycolysis (Fig. 2a). Notably, there was also a significant loss of glucose transporter 2 (*Slc2a2*) and glucokinase (*Gck*) expression, as well as up-regulation of *Ldha* and *Pdk1* (Figs. 2a,b). *Ldha* is one of several ‘disallowed’ genes, the repression of which distinguishes mature β-cells and it helps maintain optimal capacity for glucose-stimulated OXPHOS and insulin secretion^36^. The increase in *Ldha* expression was not accompanied by significant up-regulation of other core β-cell disallowed genes^37^ (Supplemental Fig. 3), but together these transcriptional changes indicate a metabolic reprogramming with increased partitioning of glucose-derived pyruvate away from mitochondria toward lactate (Fig. 2c). Of note, the HG-induced upregulation of glycolysis-related genes included *Aldh1a3,* which is a feature of dedifferentiated and metabolically deficient β-cells^38^. Closer inspection further showed higher expression of the α-cell-restricted gene *Gc*^39^, and reduced levels of key β-cell markers *Ins2, Ucn3, Mafa, Nkx6-1, Slc2a2* (*Glut2*)*, Glpr1*, and *Slc30a8*, all of which indicates a beginning loss of mature β-cell identity (Figs. 2d,e). Interestingly, *Pdx1* levels were higher in BclxβKO islets (*p*_adj_ < 0.001, BclxβKO vs BclxβWT in HG), but overall, the HG-induced transcriptional changes to glycolysis and β-cell identity were similar between the two genotypes. Together these results show that prolonged high glucose exposure causes metabolic reprogramming and beginning loss of the mature β-cell identity, and this happens similarly in BclxβWT and BclxβKO islets.

**Figure 2.**
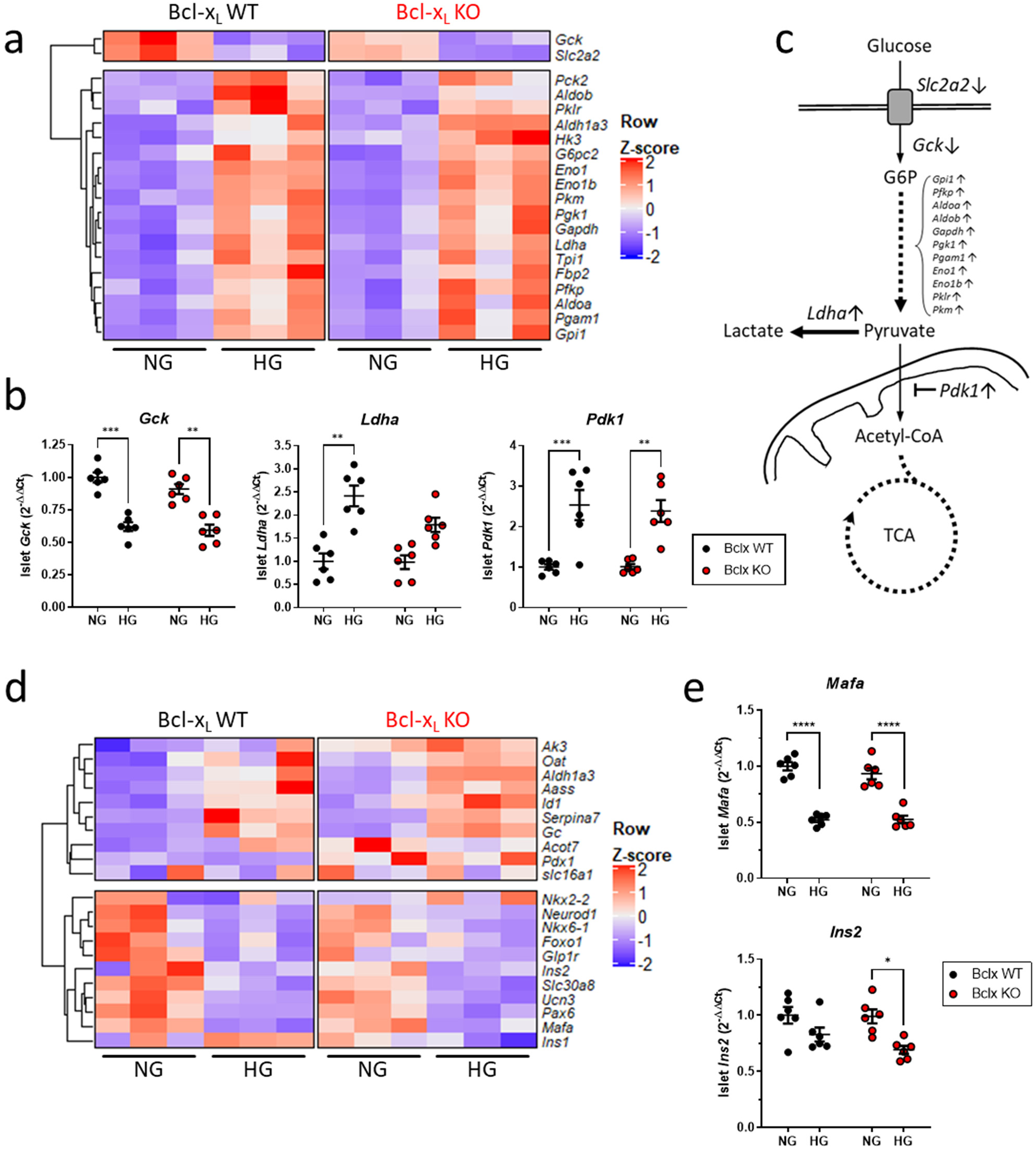
Prolonged exposure to high glucose transcriptionally re-wires β-cell glycolysis and identity. a) Heat map of differentially expressed glycolytic genes common to BclxβWT and BclxβKO islets after 6 days culture in high glucose (HG) compared to normal glucose (NG; n=3 cDNA library preparations). b) qPCR quantification of select transcripts confirming key RNA-Seq findings in panel a (n=6). c) Schematic illustrating the re-wiring of glycolysis in BclxβWT and BclxβKO islets after prolonged high glucose exposure. The transcriptional signature shows loss of β-cell genes *Slc2a2* and *Gck,* up-regulation of general glycolytic transcripts, and redirection of glucose-derived pyruvate from mitochondria toward lactate production. d) Heat map showing HG-induced changes to expression of key genes related to β-cell identity. The overall transcriptional signature is consistent with loss of the mature β-cell phenotype and beginning β-cell dedifferentiation in HG-cultured islets of both genotypes. e) qPCR confirmation of HG-induced decrease in the expression of the β-cell identity genes *Mafa* and *Ins2* (n=6). In all panels, \**p*<0.05, \*\**p*<0.01, \*\*\**p*<0.001 and \*\*\*\**p*<0.0001 by two-way ANOVA followed by Šidák’s multiple comparison test.

### Effects of excess glucose and Bcl-x_L_ deletion on β-cell Ca^2+^ signaling and insulin secretion

To determine how Bcl-x_L_ knockout and HG-induced transcriptional changes related to β-cell function, we first compared acute glucose-stimulated Ca^2+^ signalling in BclxβWT and BclxβKO islets. Voltage-gated Ca^2+^ entry is a prerequisite signal that triggers and shapes the kinetics of glucose-stimulated insulin secretion^3^. After 2 days in NG culture, BclxβWT islets responded to 15 mM glucose with a mix of Ca^2+^ oscillations (47%) and plateau rises (53%), which is expected for a glucose stimulus of intermediate strength. In contrast, all BclxβKO islets responded with plateau steady-state Ca^2+^ elevations (Fig 3a), which is normally seen at higher, saturating, glucose concentrations. After 2 days of HG culture, islets of both genotypes had modest elevations in baseline Ca^2+^ and a beginning delay in their return to basal after removal of the stimulus (Figs. 3a). Additionally, HG-cultured BclxβWT islets no longer oscillated and only showed Ca^2+^ plateaus similar to BclxβKO islets (Fig. 3a). Extending the HG culture to 6 days did not affect the plateau or peak Ca^2+^ levels in either genotype (Figs. 3b,c). However, the elevation in basal Ca^2+^ and the delay in recovering to baseline were worsened, and these disruptions to islet Ca^2+^ kinetics were exacerbated in the absence of Bcl-x_L_ (Figs. 3b,c). Ca^2+^ on- and off-rates in response to depolarization with KCl were not affected, so the inability to rapidly terminate glucose-induced Ca^2+^ signaling was not due to impaired extrusion and buffering mechanisms (Fig. 3d).

**Figure 3.**
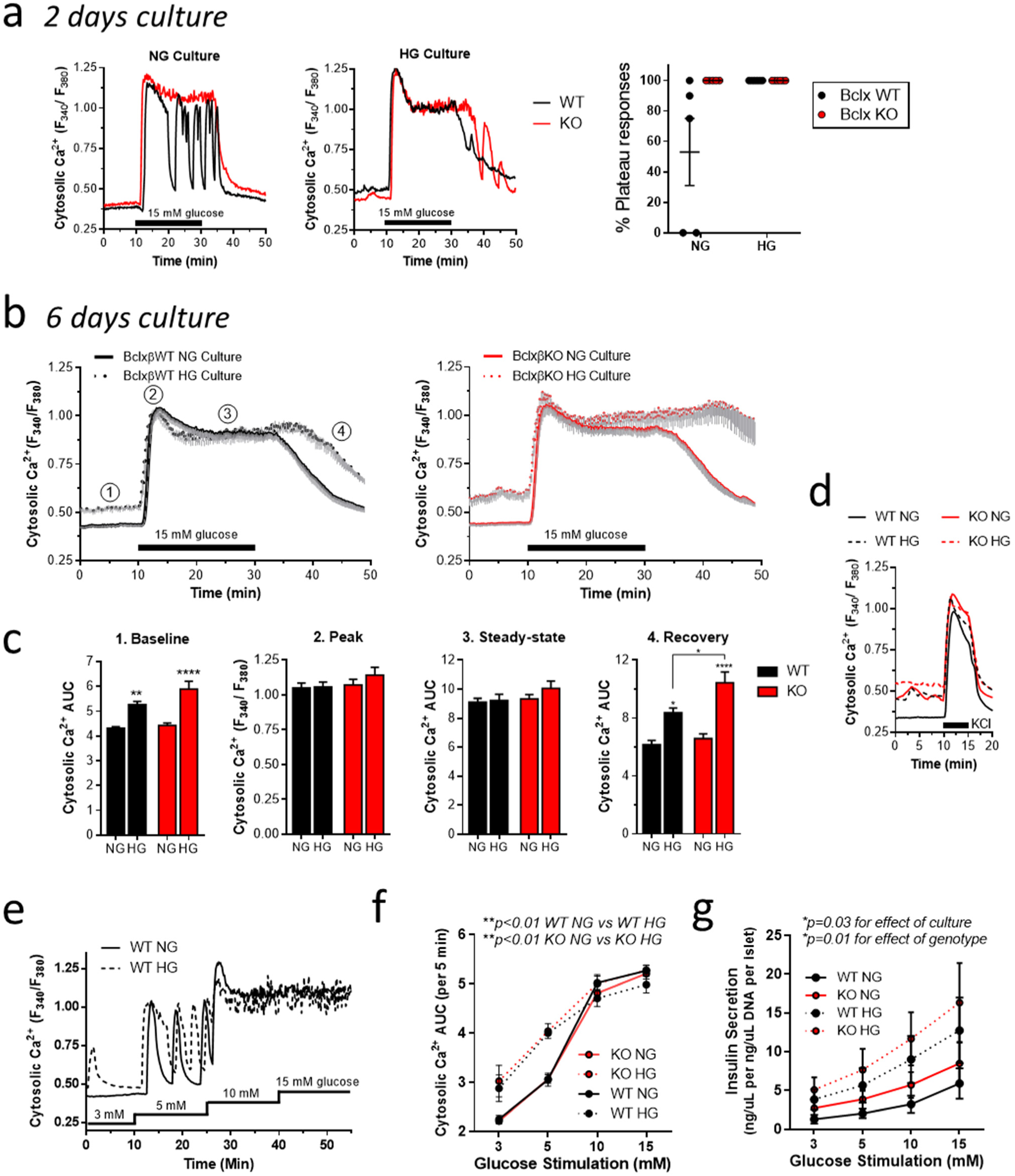
Prolonged exposure to excess glucose perturbs islet Ca^2+^ dynamics, sensitizes glucose-stimulated Ca^2+^ responses and amplifies insulin secretion. a) Representative cytosolic Ca^2+^ responses in glucose-stimulated BclxβWT and BclxβKO islets after 2d culture in 11 mM glucose (NG; *left panel*) or 25 mM glucose (HG; *middle panel*, as well as quantification of the percentage of islets showing plateau responses (*right panel*; n=5, between 1 and 14 islets in each experiment). b) Average glucose-induced cytosolic islet Ca^2+^ responses of BclxβWT and BclxβKO islets after 6 days culture in NG and HG. Gray hanging bars represent SEM (NG n=5 and HG n=3 with between 4 and 10 islets in each experiment) c) Quantification of average islet Ca^2+^ levels during the baseline (1), peak (2), steady-state (3) and recovery (4) stages of the glucose response, as indicated in panel b (NG n=5 and HG n=3, \**p*<0.05 by multiple t-tests). d) Islet cytosolic Ca^2+^ responses to metabolism-independent islet depolarization with 30 mM KCl in the presence of basal 3 mM glucose. Each trace is the averages of 4-9 islets from 3 mice of each genotype. Traces are shown without error bars to better highlight the on- and off-rates of the responses. e) Representative example of BclxβWT islet Ca^2+^ responses to step-wise glucose ramp stimulation after NG or HG culture. f) Integrated concentration-response profiles (area under the curve; AUC) demonstrating similar glucose sensitization of BclxβWT and BclxβKO islet Ca^2+^ responses after NG and HG culture (Total 70 islets from n=4 mice. \*\**p*<0.01 by two-way repeated measures ANOVA followed by Sidak’s multiple comparisons test). g) Glucose concentration-response profiles of insulin secretion from BclxβWT and BclxβKO islets after culture in NG and HG (n=6, \**p*<0.05 by two-way ANOVA of AUC values matched across each experiment, followed by Sidak’s multiple comparisons test).

A transition from oscillatory to plateau Ca^2+^ responses at intermediate glucose levels may occur if islets are sensitized to the sugar. We therefore tested the responsiveness of NG- and HG-cultured islets by comparing cytosolic Ca^2+^ during a step-wise glucose ramp, exemplified in Fig. 3e. As quantified in Fig. 3f, HG pre-culture left-shifted the glucose-response profiles of BclxβWT and BclxβKO islets to a similar degree. Importantly, HG-cultured islets also showed amplified insulin secretion across the same range of glucose concentrations (Fig. 3g). Although the two genotypes had similarly sensitized Ca^2+^ responses, BclxβKO islets secreted significantly more insulin than BclxβWT islets after both NG and HG culture. Islet insulin content was measured after the secretion assay and did not differ by genotype. The contents were decreased in HG cultured islets, but this was accounted for by the amplified insulin release during the glucose ramp (Supplemental Fig. 4).

Collectively, our data thus far show that islets respond to prolonged glucose excess with compensatory sensitization of Ca^2+^ responses and amplified insulin secretion, despite transcriptional rewiring of glycolysis and the onset of β-cell dedifferentiation. At this stage, islet Ca^2+^ response kinetics are perturbed, and both Ca^2+^ dysregulation and insulin hyper-secretion are exacerbated in the absence of β-cell Bcl-x_L_.

Bcl-x_L_ is needed to preserve mitochondrial transcripts and mitochondrial hyperpolarization in β-cells under high glucose culture

To better understand the specific role of Bcl-x_L_ in the β-cell response to chronic glucose excess, we next focused on the genotype-dependent differences in gene expression. In NG culture, a total of 52 genes were differentially expressed between BclxβWT and BclxβKO islets. High glucose exposure caused a substantial divergence of the transcriptional profiles and the number of differentially expressed genes increased to 234 (Fig. 4a). Only 4 of the differentially expressed transcripts were common to the two culture conditions, including *Bcl2l1,* which encodes for Bcl-x_L_ and *Esr1,* which we believe reflects the expression of the CreER^TM^ transgene in BclxβKO islets. Functional annotation and enrichment analysis showed that the genotype-dependent differences following HG culture were dominated by transcripts related to mitochondria (Fig. 4b). In total, 53 genes fell in the top Cell Component GO-term “Mitochondrion” and 50 (94%) of these were down-regulated in BclxβKO islets compared to BclxβWT. Notably, a large number of these genes encode for subunits of electron transport chain (ETC) complexes I through IV, as well as subunits of the ATP Synthase (Complex V) and mitochondrial ribosomal proteins (Fig. 4c). We confirmed the striking culture- and genotype-dependent differences in key OXPHOS transcripts by qPCR (Fig. 4d).

**Figure 4.**
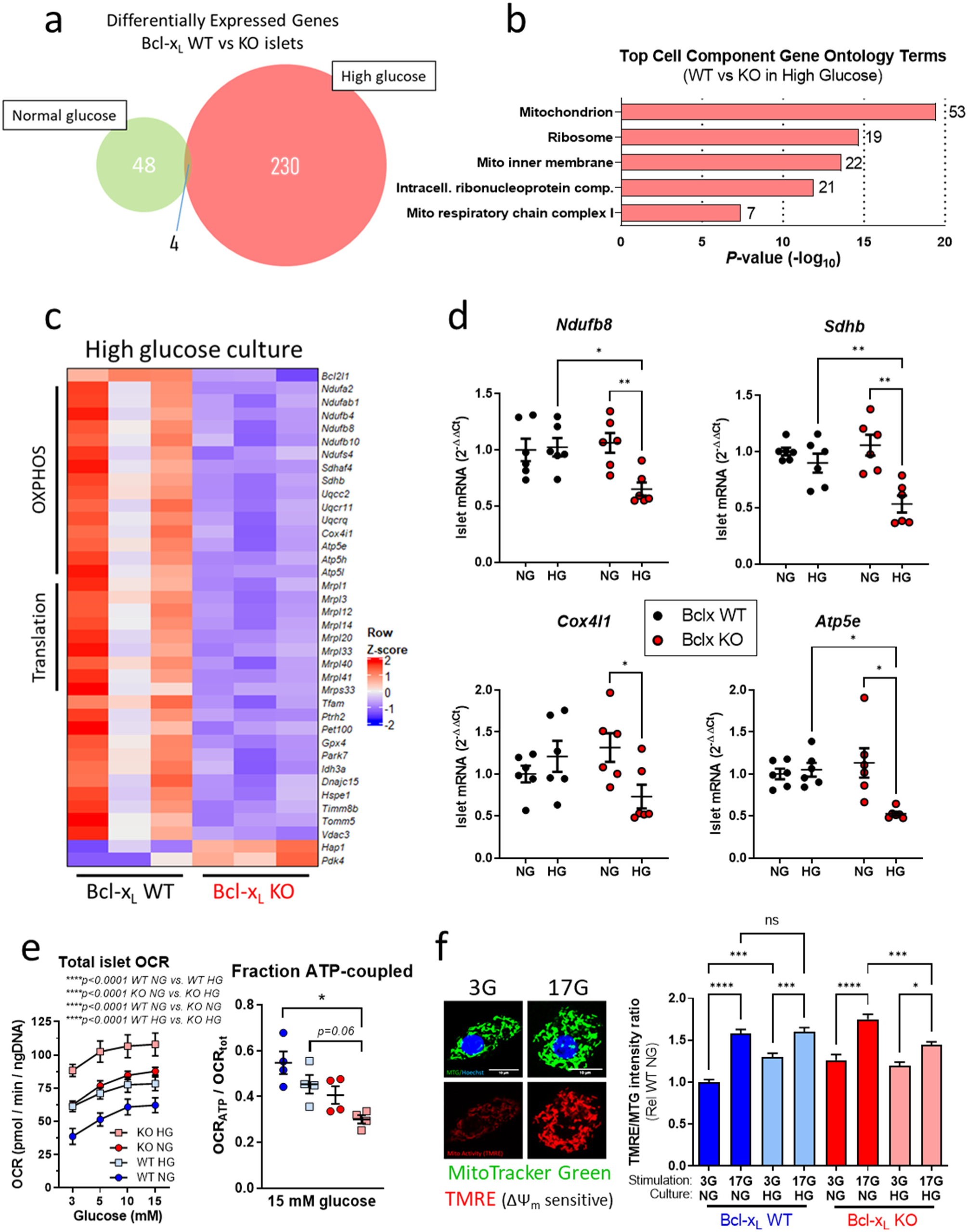
Bcl-x_L_ preserves the expression of mitochondrial transcripts and glucose-stimulated mitochondrial hyperpolarization during high glucose-stress. a) Venn diagram of differentially expressed genes between BclxβWT and BclxβKO islets after 6 days culture in normal glucose (11 mM; NG) or high glucose (25 mM; HG). b) Top cell component GO terms identified by enrichment analysis and functional annotation of the differentially expressed genes between HG-cultured BclxβWT and BclxβKO islets. The number of transcripts in each category is indicated by the bars. c) Heat map of select differentially expressed genes from the top CC GO-category “Mitochondrion” in panel b. d) qPCR quantification of select OXPHOS-related transcripts confirming key RNA-Seq findings in panels a-c (n=6, \**p*<0.05, \*\**p*<0.01 by two-way ANOVA followed by Tukey’s multiple comparisons test). e) *Left* - Total oxygen consumption rates (OCR_tot_) of NG- and HG-cultured BclxβWT and BclxβKO islets in response to stimulation with progressive increases in extracellular glucose (n=6; \*\**p*<0.01, \*\*\**p*<0.001 by Two-way ANOVA followed by Tukey’s multiple comparisons test). *Right* – Fraction of total OCR that is coupled to ATP synthesis in the presence of 15 mM glucose (n=4, *\*\*p*<0.01 by two-way ANOVA followed by Tukey’s multiple comparisons test). f) *Left* – Examples of β-cells double-stained with MitoTracker Green (MTG; ΔΨ_m_-insensitive) and TMRE (ΔΨ_m_-sensitive) in the presence of basal 3 mM glucose (3G) and stimulatory 17 mM glucose (17G). *Right –* Quantification of the effects of acute glucose stimulation on ΔΨ_m_ (TMRE/MTG ratio) in NG- and HG-cultured cells (87 to 91 cells in each group from 7 mice of each genotype; \**p*<0.05, \*\*\**p*<0.001, \*\*\*\**p*<0.0001 by one-way ANOVA followed by Sidak’s multiple comparisons test).

We next compared mitochondrial function by measuring glucose-stimulated OCR and mitochondrial membrane potential (ΔΨ_m_). Pre-culture in high glucose significantly amplified total OCR (OCR_tot_) in islets from both genotypes across a range of glucose stimuli. Under all conditions, BclxβKO islets consumed significantly more oxygen than BclxβWT islets (Fig. 4e). In agreement with the increased basal OCR, HG pre-culture also increased the basal ΔΨ_m_ of dispersed BclxβWT β-cells in 3 mM glucose, but the ΔΨ_m_ response to acute stimulation by glucose was not amplified (Fig. 4f). Notably, BclxβKO cells developed a defect in glucose-stimulated ΔΨ_m_ hyperpolarization when cultured in HG (Fig. 4f). This suggests Bcl-x_L_ helps β-cells preserve their mitochondrial proton-motive force when challenged by chronic glucose excess. By comparing OCR in the presence and absence of the ATP Synthase inhibitor oligomycin, we quantified the amount of glucose-stimulated oxygen consumption that was coupled to ATP synthesis (OCR_ATP_)^18,40^. Despite the drastic increase in total OCR, HG culture did not increase OCR_ATP_ in either genotype. Consequently, HG notably reduced the fraction of OCR_tot_ that was coupled to ATP production, particularly in BclxβKO islets (Fig. 4e). This indicates that the impaired ΔΨ_m_ response in Bcl-x_L_ KO β-cells is a consequence of respiratory uncoupling.

### Bcl-x_L_ is required for normal β-cell expression of Tfam and Pgc-1α

Our RNA-Seq analysis (Fig. 4c) showed an intriguing loss of the mitochondrial transcription factor *Tfam* specifically in HG-cultured BclxβKO islets. Tfam is essential for transcription and replication of the mitochondrial genome, as well as packaging of mitochondrial DNA (mtDNA) into nucleoids^41^. In β-cells, Tfam is important for mitochondrial function and insulin secretion^42,43^. We therefore confirmed the BclxβKO-specific loss of *Tfam* by qPCR (Fig. 5a). *Tfam* levels are transcriptionally regulated by Nrf-1 and its co-regulator Pgc-1α, which also control the expression of a large number of nuclear genes involved in mitochondrial biogenesis and function, including OXPHOS components. We examined *Ppargc1a* expression and found that Pgc-1α mRNA levels were increased by HG-culture in BclxβWT islets. In marked contrast, the induction of Pgc-1α transcript by glucose stress was completely absent in BclxβKO islets and β-cells (Figs. 5a). The dysregulation of Pgc-1α and Tfam expression in BclxβKO β-cells suggests Bcl-x_L_ may also be important for the control of mitochondrial mass.

**Figure 5.**
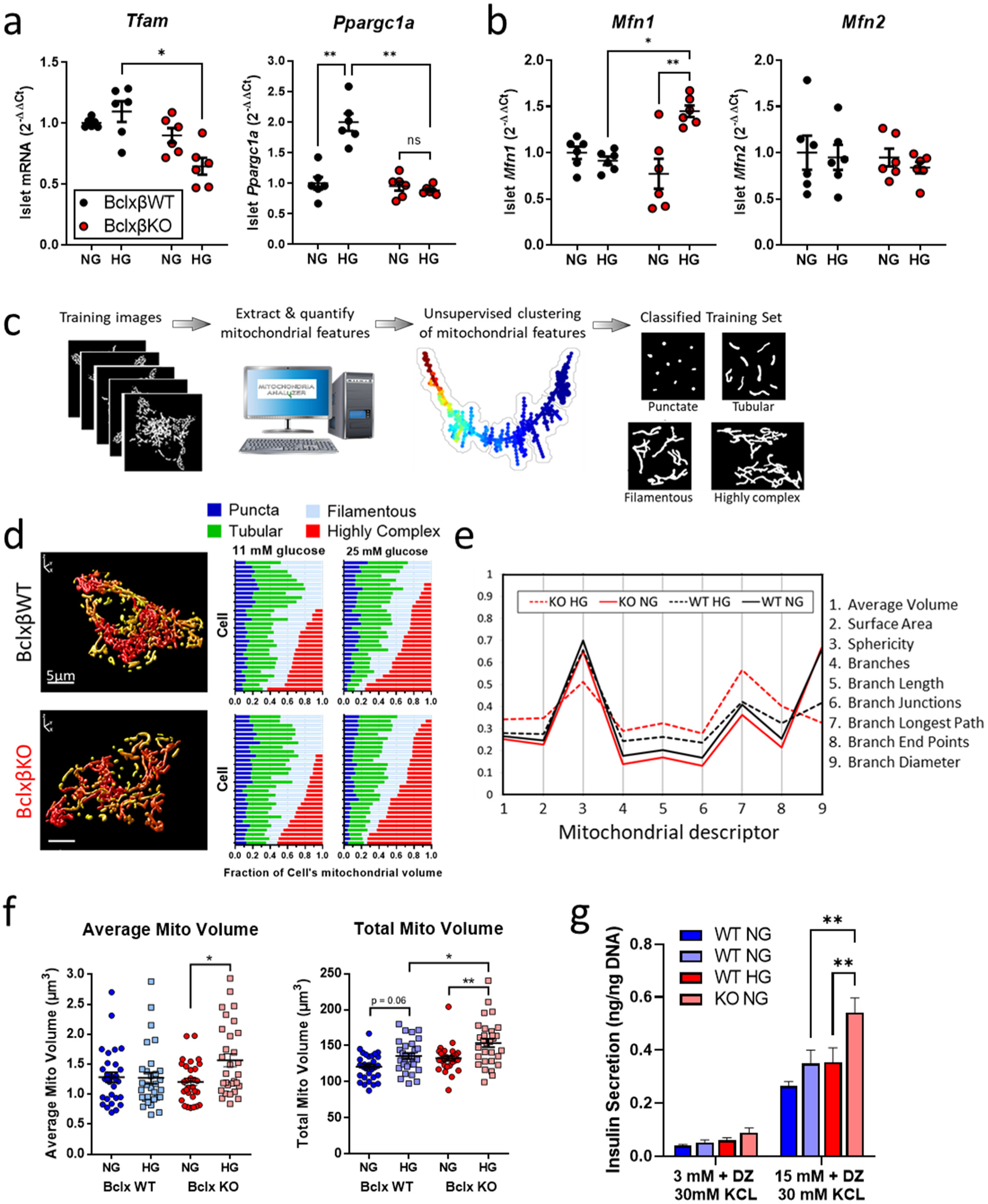
Bcl-x_L_ KO β-cells respond to prolonged high glucose with higher mitochondrial network complexity, increased mitochondrial volume, and augmented insulin secretion in the amplifying pathway. a) qPCR quantification of *Tfam* and *Pgc-1α* mRNA in BclxβWT and BclxβKO islets following 6 days culture in NG and HG (n=6). b) qPCR analysis of the expression of key mediators of mitochondrial fusion (*Mfn1, Mfn2*) in BclxβWT and BclxβKO islets after 6 days culture in NG or HG (n=6). c) Schematic illustration of workflow for generation of training set for machine learning-based classification of 3D mitochondrial morphometric features. d) *Left:* Examples of 3D confocal reconstruction of mitochondrial networks in Bcl-x_L_ WT and Bcl-x_L_ KO cells. *Right*: Machine learning-based classification of mitochondrial morphology distributions in Bcl-x_L_WT and Bcl-x_L_ KO cells after NG or HG culture. (3 mice of each genotype. 30 cells were examined from each mouse and a total of 120 cells and ∼16,000 individual were mitochondria classified). e) Parallel coordinates plot comparing normalized means of all extracted mitochondrial morphometric and network complexity features across genotypes and culture conditions. f) Quantification of average mitochondrial volume and total mitochondrial volume in NG- and HG-cultured cells (n=30 cells per genotype in each condition). g) Comparison of the amplifying pathway of insulin secretion in NG- and HG-cultured BclxβWT and BclxβKO islets. Plasma membrane potential was uncoupled from β-cell glucose metabolism by clamping ATP-sensitive K^+^-channels open with 250 µM diazoxide (DZ). Insulin secretion was then measured following direct depolarization by 30 mM KCl in the presence of 3 mM glucose or 15 mM glucose (n=6). In all panels, *\*p*<0.05, \**\*p*<0.01 by two-way ANOVA followed by multiple comparisons tests.

### Bcl-x_L_ limits high glucose-induced perturbations of β-cell mitochondrial dynamics and mass

Mitochondrial function, and their adaptation to metabolic demand, are closely interrelated with the regulation of mitochondrial network morphology and total mitochondrial mass. Network connectivity is shaped by the balance of dynamic fusion and fission events^44,45^, while mass is dictated by the relative amounts of mitochondrial biogenesis versus degradation by macroautophagy and mitophagy^46,47^. The effects of prolonged glucose excess on β-cell mitochondrial morphofunction and abundance, however, remain unclear. It is also an important unanswered question if Bcl-x_L_ affects mitochondrial dynamics in β-cells, as reported in neurons^48^.

HG culture significantly increased expression of the mitochondrial fusion regulator *Mfn1* exclusively in BclxβKO islets. The mRNA levels of other fusion-mediators *Mfn2* and *Opa1*, as well as the mitochondrial fission regulators *Drp1* and *Fis1,* were not significantly affected by HG challenge in either genotype (Fig. 5b and Supplemental Fig. 5). To determine the overall impact on mitochondrial networking, we used our recent pipeline for confocal analysis of mitochondrial dynamics and the associated Mitochondria Analyzer software tool^19,20^. For the most accurate comparisons^19,21^, we generated full 3D reconstructions of the mitochondrial networks and extracted 9 parameters to quantitatively describe the morphology and structural complexity of each individual organelle. In parallel, we performed hierarchical clustering of parameters extracted from independent imaging experiments, which identified 4 primary morphological categories of β-cell mitochondria; designated as Puncta, Tubular, Filamentous, and Highly Complex (see Table 1 in Methods for defining features). We then established a training set and performed an unbiased classification of ∼16,000 individual mitochondria from NG- and HG-cultured BclxβWT and BclxβKO cells into the 4 morphological categories using machine learning algorithms (Fig. 5c and Methods). By calculating the fraction of each cell’s total mitochondrial volume that was mapped to each category, we found that HG culture significantly increased the population of Highly Complex mitochondria in BclxβKO cells (*p*<0.005, HG vs NG), but not in BclxβWT cells (*p*=0.36; HG vs NG) (Fig. 5d). Combined with the increase in *Mfn1*, this indicates an increase in mitochondrial fusion, which was substantiated by a larger average organelle volume, quantified on a per-cell basis (Fig. 5f). Using a parallel coordinates plot we can get an overview of how mitochondrial parameters vary between genotype and culture conditions (Fig. 5e), and this highlights that BclxβKO cells respond to high glucose with the most pronounced changes to all descriptors of mitochondrial morphology and connectivity. Importantly, BclxβKO cells also had a marked increase in total mitochondrial volume that was not seen in BclxβWT cells (Fig. 5f). Because mitochondrial branch diameter and sphericity were reduced (Fig. 5e), it is unlikely that the additional volume reflects stress-induced damage and swelling.

The observed increase in total mitochondrial volume might explain how HG-cultured BclxβKO islets remain glucose-responsive and hyper-secrete insulin, despite a significant loss of mitochondrial transcripts and ΔΨ_m_. Our RNA-Seq analyses did not indicate major differences in TCA cycle components, suggesting the defects might be predominantly ETC-related. If so, an increase in mitochondrial volume should still enhance the formation of mitochondria-derived factors for amplification of insulin secretion^49^. Indeed, glucose-dependent amplification of KCl-stimulated insulin secretion in the presence of diazoxide was significantly augmented in BclxβKO islets, compared to BclxβWT (Fig. 5g), consistent with a volume-dependent compensation for ETC-specific defects.

### Mitochondrial ROS drives transcriptional reprogramming and Bcl-x_L-_dependent differences in mitochondrial homeostasis

We previously reported that Bcl-2 has functions in β-cell redox control, and that small molecule co-inhibitors of Bcl-2 and Bcl-x_L_ promote β-cell ROS formation^11^. A specific role of Bcl-x_L_ was not examined, but ROS could conceivably link Bcl-x_L_ to changes in mitochondrial dynamics and function^50^, as well as mitochondrial transcription and retrograde control of nuclear gene expression^47,51–53^.

We examined mitochondrial ROS (mitoROS) levels using the superoxide-specific sensor MitoSOX and found that deletion of β-cell Bcl-x_L_ caused a significant mitoROS increase in NG cultured cells, and that HG culture amplified mitoROS in cells of both genotypes (Fig. 6a). MitoROS levels could be normalized using the mitochondria-targeted anti-oxidant MitoTEMPO (Fig. 6a), which substantiates that the mitoROS is mitochondrial and likely ETC-derived. To determine the metabolic consequences of the observed ROS differences, we cultured BclxβWT and BclxβKO islets in NG or in HG with and without the addition of MitoTEMPO and then exposed them to a mitochondrial stress test, which incorporated an acute stimulation with 15 mM glucose (Fig. 6b). As expected, HG pre-culture elevated basal and glucose-stimulated OCR (Fig. 6c). Respiratory control analysis^18^ substantiated our previous observation that β-cell Bcl-x_L_ deficiency and HG culture impair ATP coupling efficiency, and further showed that this was associated with the development of a major mitochondrial proton leak (Fig. 6d). Consistent with the observed increases in mitochondrial mass (Fig. 5f), Bcl-x_L_ deletion and HG culture also notably amplified spare respiratory capacity. These perturbations to islet respiratory parameters were all mitigated by lowering of mitoROS levels (Fig. 6d). Scavenging mitoROS during HG culture also amplified the already elevated Ca^2+^ responses to stimulation with low glucose concentrations (Fig. 6e). ROS has roles in physiological signal transduction, but this result indicates that mitoROS limits, rather than mediates, sensitization of the triggering Ca^2+^ signal for glucose-stimulated insulin secretion in HG-cultured islets.

**Figure 6.**
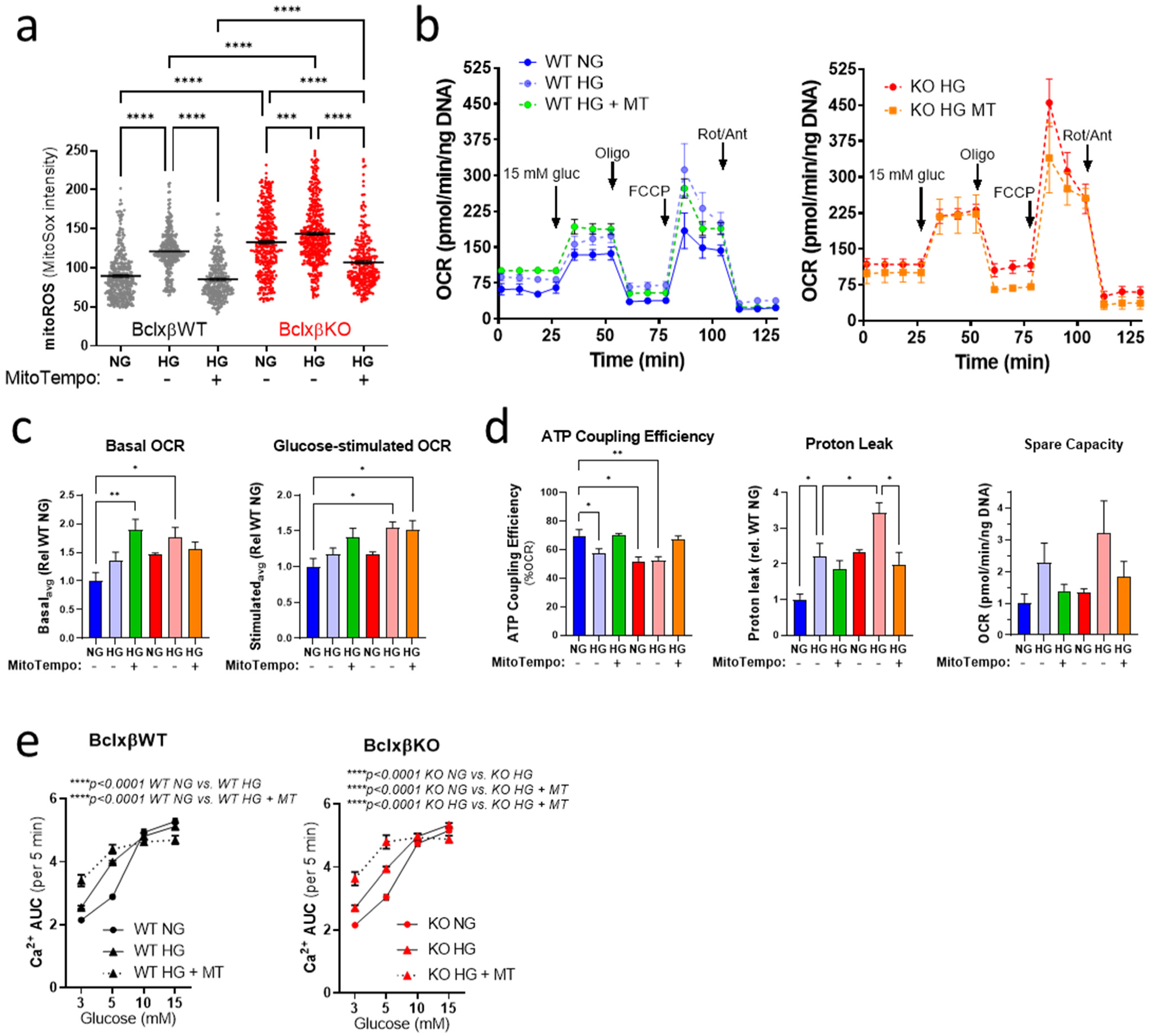
Bcl-x_L_-dependent regulation of mitoROS preserves ATP-coupled islet respiration and limits the sensitization of basal Ca^2+^ responsiveness during high glucose-stress. a) Mitochondrial superoxide (mitoROS) levels imaged by MitoSox staining in BclxβWT and BclxβKO islet cells after 6 days culture in NG or HG with and without the addition of 10 µM mitoTEMPO (276 to 427 cells in each group from 2 independent preparations). b) Example of Seahorse mitochondrial stress tests on BclxβWT and BclxβKO islets after HG culture with and without the addition of mitoTEMPO (MT). At the indicated time-points wells were injected with 15 mM glucose, 1 µM oligomycin (Oligo), 2 µM FCCP, and a combination of 1 µM Rotenone and 1 µM antimycin A (Rot/Ant). For a clearer comparison, NG-cultured islet traces are not shown, but are included in the quantifications in panel c. c) Respiratory control analysis based on mitochondrial stress tests of BclxβWT and BclxβKO islets after culture in NG, or in HG with and without mitoTEMPO (n=3-5 islet preparations in each group). d) Integrated Ca^2+^ responses (area under the curve; AUC) to stimulation by a step-wise glucose ramp. Responses of BclxβWT and BclxβKO islets were measured after 6 days NG culture, and after HG culture with and without the addition of mitoTEMPO (MT; n=3, 4-9 islets per experiment). In panels a & c \**p*<0.05, \*\**p*<0.01, \*\*\**p*<0.001, \*\*\*\**p*<0.0001 by one-way ANOVA followed by multiple comparison tests. In panel d, \*\*\*\**p*<0.0001 by two-way ANOVA followed by Tukey’s multiple comparisons test.

We next assessed the role of mitoROS in the HG-induced changes to β-cell gene expression. Remarkably, all major glucose-induced transcriptional changes we assessed by qPCR were partially or fully reversed by normalization of mitochondrial ROS (Fig. 7a,b and Supplemental Fig. 6). This included rescue of the HG-induced changes related to glycolytic flux and β-cell identity (*Gck*, *Ldha*, *Pdk1, Ins2, Mafa*) that were common to both genotypes, as well as the loss of *Tfam*, and ETC components (*Ndufb8*, *Sdhb*, *Cox4l1, Atp5e*) that was specific to BclxβKO islets.

**Figure 7.**
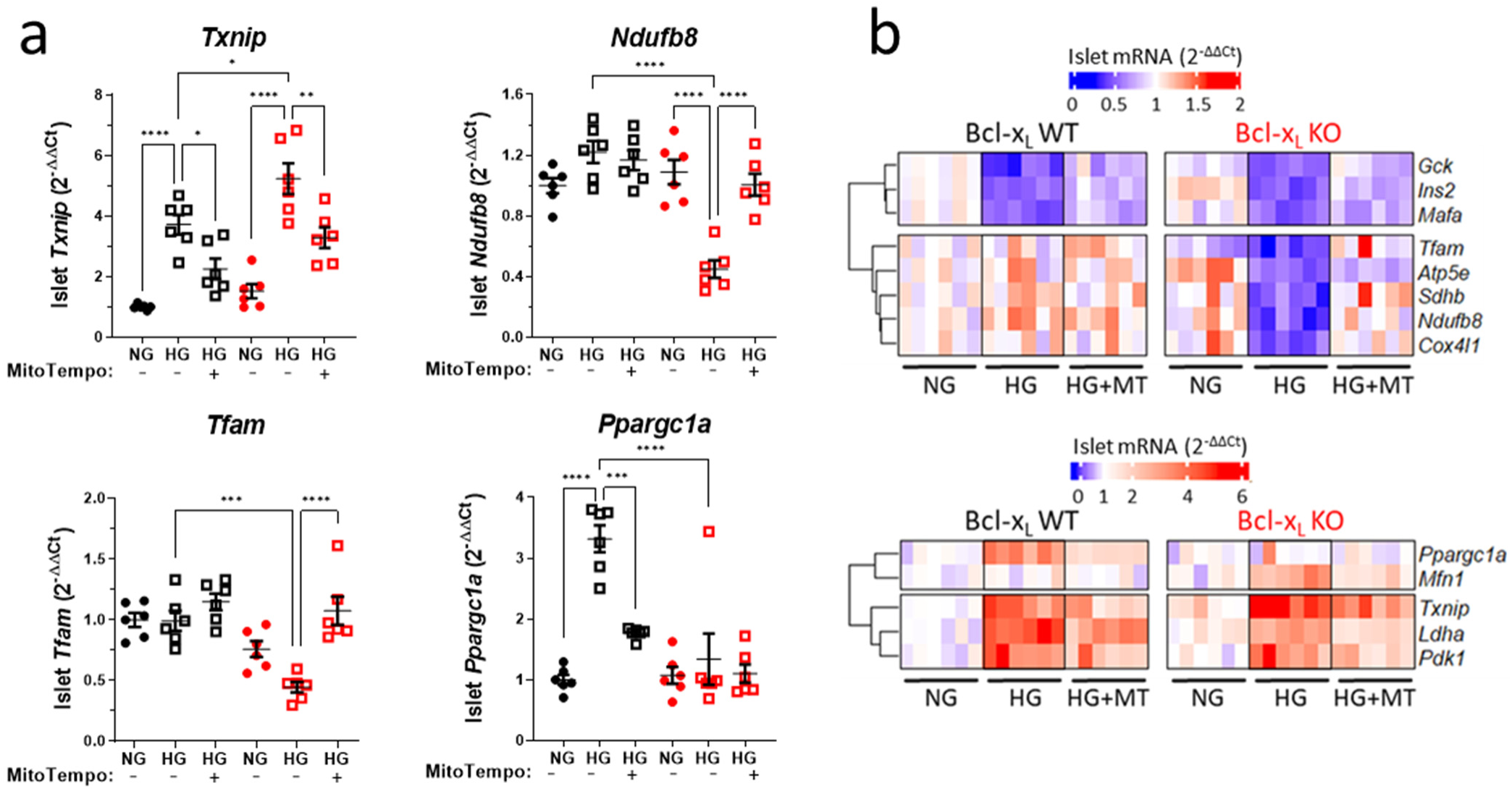
High glucose-induced transcriptional dysregulation in Bclx_L_-deficient β-cells is driven by mitoROS. a) qPCR quantification of select mRNAs in BclxβWT and BclxβKO islets following 6 days culture in NG and HG with and without the addition of 10 µM mitoTEMPO (n=6; \**p*<0.05, \*\**p*<0.01, \*\*\**p*<0.001, \*\*\*\**p*<0.0001 by one-way ANOVA followed by Tukey’s multiple comparison tests). Heat map summarizing qPCR quantification of islet mRNAs after culture in NG, or in HG with and without mitoTempo (MT). *Top:* transcripts that were down-regulated in HG. *Bottom:* transcripts that were up-regulated in HG. All corresponding qPCR data are shown in panel a and Supplemental Figure 5.

## DISCUSSION

In this study, we characterized the pancreatic β-cell response to sub-lethal levels of glucose stress, and used conditional gene deletion to determine the role of the anti-apoptotic protein Bcl-x_L_ in these processes. In both wild-type and Bcl-x_L_ knockout islets, prolonged glucose excess caused a robust transcriptional reprogramming of metabolism toward increased glycolysis and beginning loss of the mature β-cell identity. These changes to gene expression in large part resemble the signatures of hyperglycemia-induced β-cell dedifferentiation in partially pancreatectomized rats^54,55^. Interestingly, our results showed that in glucose-stressed cells, the transcriptional loss of β-cell identity occurs while there is still compensatory β-cell hyper-function, and precedes the development of maladaptive ER stress. This indicates that loss of the mature differentiated β-cell phenotype manifests remarkably early in the progression of glucotoxicity. Whether this sequence of events differs under other forms of diabetogenic stress is not clear.

Maladaptive ER stress, has been proposed as a primary event preceding oxidative and inflammatory stress in T2D^30^. By all indications, the glucose-induced changes to β-cell UPR progressed similarly in BclxβWT and BclxβKO islets, suggesting that Bcl-x_L_ does not modulate the susceptibility of β-cells to ER stress, unlike the pro-apoptotic Bax and Bak^56,57^. Rather, we identified a mitochondria-specific defect in the absence of Bcl-x_L_. Prolonged high-glucose exposure amplified respiration, but Bcl-x_L_-deficient β-cells developed mitochondrial dysfunction with reduced coupling of ETC flux to ΔΨ_m_ hyperpolarization and ATP synthesis. Importantly, our results also revealed that Bcl-x_L_ is essential for normal β-cell transcription of OXPHOS components and mitochondrial ribosomes under chronic glucose excess. Overall, the loss of mitochondrial transcripts in glucose-stressed BclxβKO islets is similar to the metabolic gene expression signatures seen 2 weeks after the onset of hyperglycemia in non-obese diabetic mice^6^, and during late stages of diabetes progression in the non-obese GK rat^8^. BclxβKO islets also showed an exacerbated perturbation of basal and glucose-stimulated Ca^2+^ response kinetics. Together, this suggests that as glucose levels increase, Bcl-x_L_ counteracts mitochondrial decompensation, and thereby likely the development of a detrimental feed-forward loop of β-cell dysfunction and worsening hyperglycemia.

It is noteworthy that BclxβKO islets remained hyper-responsive despite marked loss of mitochondria-related gene expression and beginning uncoupling of respiration from ΔΨ_m._ The ability of BclxβKO islets to uphold Ca^2+^ and secretory responses in the face of beginning mitochondrial dysregulation was associated with profound changes to mitochondrial morphology and volume. Previous work has shown that mitochondrial fusion serves as a protective response to sustain survival under nutrient stress^58^. Our results indicate that mitochondrial fusion and expansion of total mitochondrial mass also are integral to functional compensation of β-cells, prior to the failure and mitochondrial fragmentation that characterizes β-cells in rodents and humans with established T2D^4–6^. In HG-cultured Bcl-x_L_ KO cells, the total mitochondrial volume increased despite the absence of glucose-induced *Ppargc1a* expression, which may reflect the presence of Pgc-1α-independent mechanisms for maintenance of mitochondrial mass in β-cells^59^. Alternatively, mitochondrial hyper-fusion in Bcl-x_L_ KO β-cells might contribute to the expansion of total mass by counteracting mitochondrial degradation by autophagy and mitophagy^60^.

Using a mitochondria-targeted antioxidant, we identified sub-lethal formation of mitoROS as a primary driver of transcriptional changes to both glycolysis and β-cell identity in glucose-stressed islets. In both NG and HG culture, Bcl-x_L_-deficient β-cells had higher mitoROS levels than wild-type. Importantly, mitochondrial antioxidants normalized respiratory coupling and restored nuclear-encoded mitochondrial gene expression in HG-cultured BclxβKO islets. This indicates that Bcl-x_L,_ protects mitochondrial homeostasis in glucose-stressed β-cells by limiting the levels of mitoROS. We previously reported that Bcl-2 dampens the physiological formation of peroxides in β-cells^11^, but a similar role for Bcl-x_L_ was not examined. The excess mitoROS in Bcl-xL KO β-cells is likely produced in the ETC as a consequence of the overall increase in mitochondrial respiration. It remains to be established how deletion of Bcl-x_L_ accelerates β-cell metabolic activity and insulin secretion. The transcriptional profiles of BclxβKO and BclxβKO islets in normal culture did not indicate differences in major metabolic pathways, so changes to protein-protein interactions likely play a role. In neurons, Bcl-x_L_ increases metabolic efficiency by limiting an ion leak conductance through the F_1_F_o_ ATPase complex^61^. Loss of such a stabilizing function may well contribute to proton leak and respiratory uncoupling in Bcl-x_L_ KO β-cells, but it would be expected to impair glucose-stimulated Ca^2+^ entry, insulin secretion and mitoROS production, in contrast to what we observe. Additional mechanisms are therefore likely at play, and future metabolomics profiling may help clarify the functions of Bcl-x_L_ in β-cell energetics.

In summary, we have identified important functions for Bcl-x_L_ in preserving mitochondrial integrity in pancreatic β-cells under non-apoptotic levels of metabolic stress. Our findings provide new insights into non-apoptotic roles Bcl-2 family survival proteins in the control of organelle physiology. We further revealed central role for mitoROS in both the functional and transcriptional impact of chronic glucose excess on β-cells. Future work is warranted to clarify the relationship between Bcl-x_L,_ mitoROS, and mitochondrial homeostasis in the pathogenesis and treatment of T2D.

## ACKNOWLEDGEMENTS

This work was supported by an Operating Grant to D.S.L. from the Canadian Institutes for Health Research (CIHR; MOP-119537). D.S.L. was supported by JDRF (CDA 2-2013-50) and BC Children’s Hospital Research Institute (BCCHR). R.S. was supported by a Canada Graduate Scholarship from CIHR. We acknowledge Dr. Jingsong Wang (BCCHR Imaging Core), as well as Mei Tang and Mitsuhiro Komba, (BCCHR) for expert technical assistance.

## AUTHOR CONTRIBUTIONS

D.J.P. and R.S. contributed equally to this work. D.J.P., R.S. and D.S.L. designed the study and wrote the paper. D.J.P., R.S., B.V., A.Z.L.S., Y.Z., A.C. and D.S.L. performed experiments and analyzed data. D.J.P., R.S., B.V., A.Z.L.S., Y.Z., A.C., B.G.H. and D.S.L interpreted data and edited the paper.

---

**Supplemental Figure 1.**
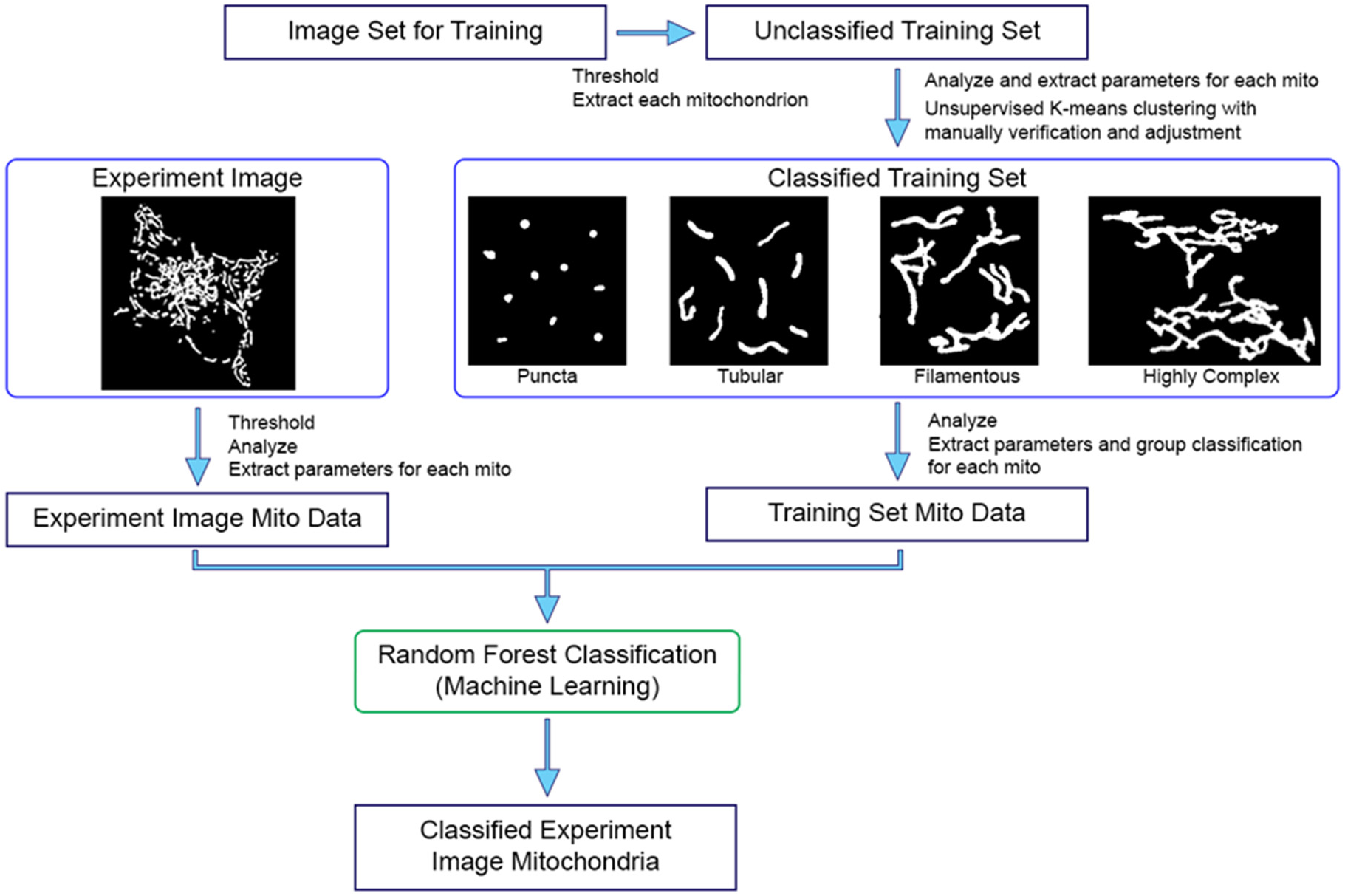
Schematic of workflow for machine learning-based classification of mitochondria. Morphometric and network connectivity features are extracted from 3D reconstructions of β-cell mitochondria and the features grouped by unsupervised K-means clustering. The initially identified groups (7 in this case) are then concatenated and manually inspected, revealing 4 major mitochondrial classifications and training set is established. Representative images from each classification are shown here as maximum projections of their original 3D object. The training set is then used for Random Forest of mitochondria that have been analyzed in independent experiments, and each mitochondrion is assigned a morphometric classification. K-means clustering and Random Forest Classification were carried out using XLSTAT.

**Supplemental Figure 2 (related to Fig. 1).**
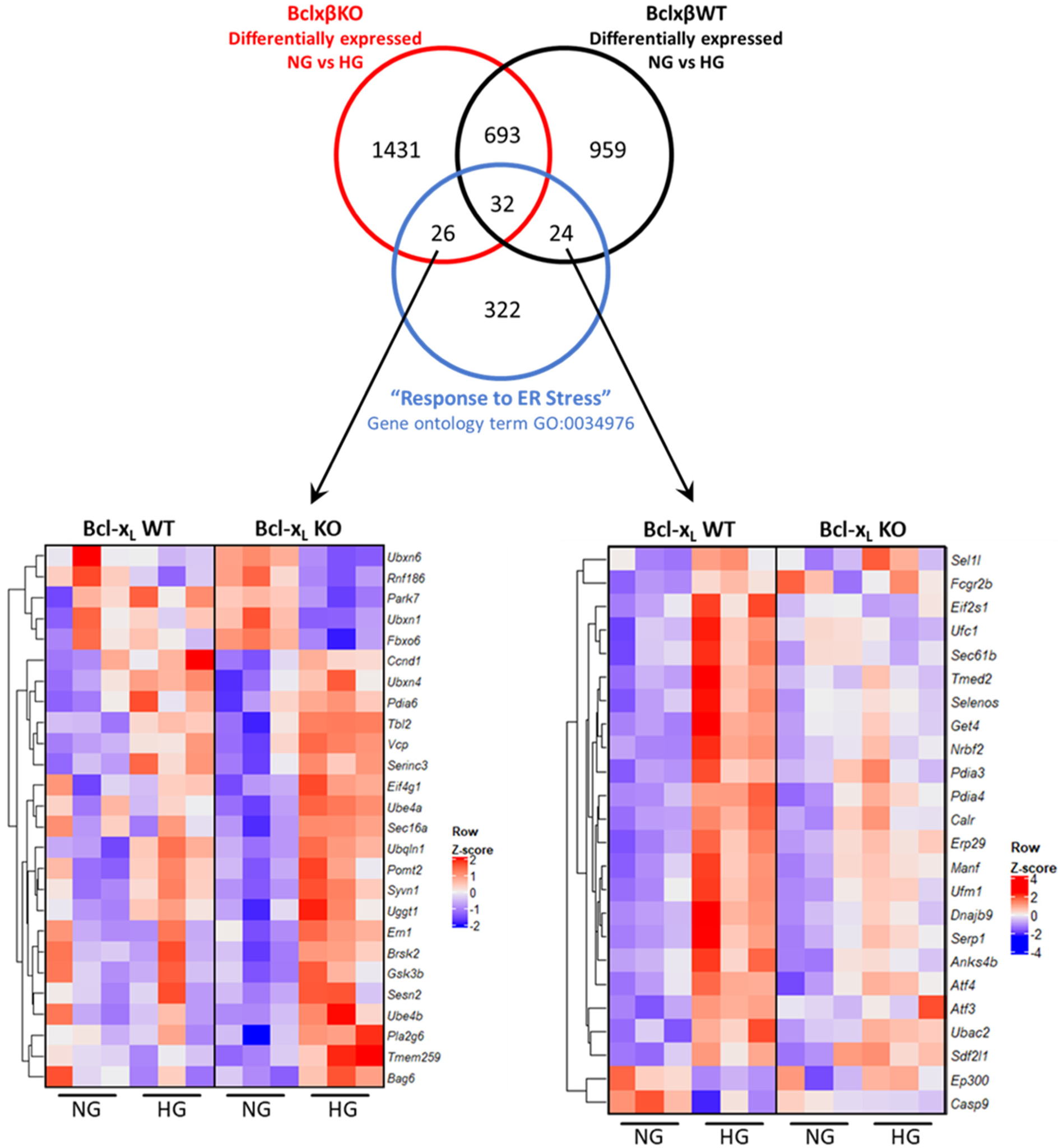
Genotype-specific changes to ER-stress-related transcripts by high glucose stress. Venn diagram and heat map showing ER stress-related genes that changed significantly in only BclxβWT islets, or only in BclxβKO islets after 6 days HG culture. The 32 common transcripts that overlap with the Gene Ontology te1m “Cellular response to ER stress” (GO:003497) are shown in Fig. 1.

**Supplemental Figure 3 (related to Fig. 2).**
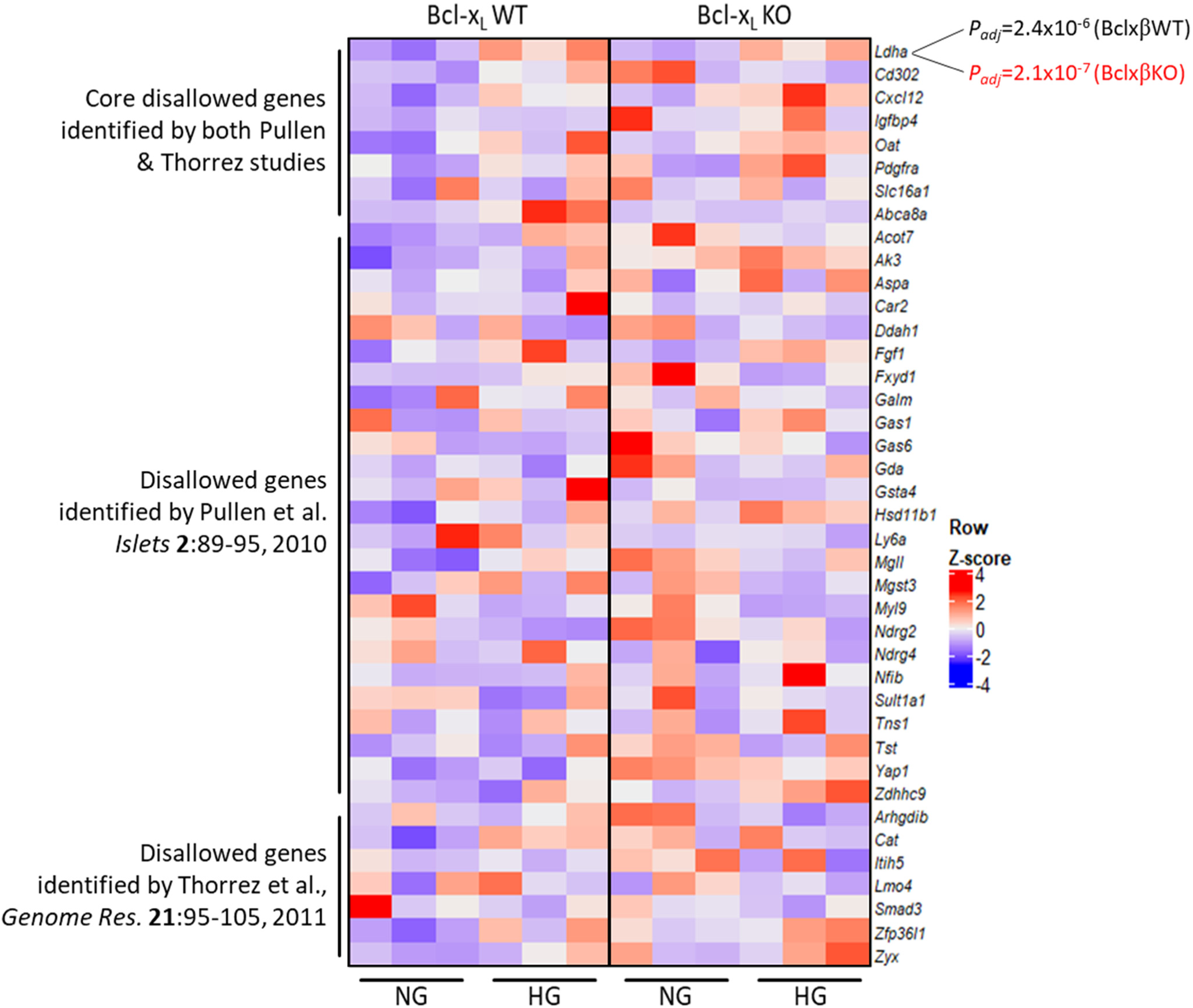
Expression of β-cell disallowed genes. Heat map showing the levels of β-cell ‘disallowed genes’ in NG- and HG-cultured BclxβWT and BclxβKO islets. The list is combines the β-cell disallowed genes identified in Pullen et al. *Islets* 2:89­95, 2010 and in Thorrez et al., *Genome* Res. 21:95-105, 2011. Of the “Core 7” shared between the two studies, only *Ldha* was significantly altered by HG culture in our RNA-Seq analysis.

**Supplemental Figure 4 (related to Fig. 3).**
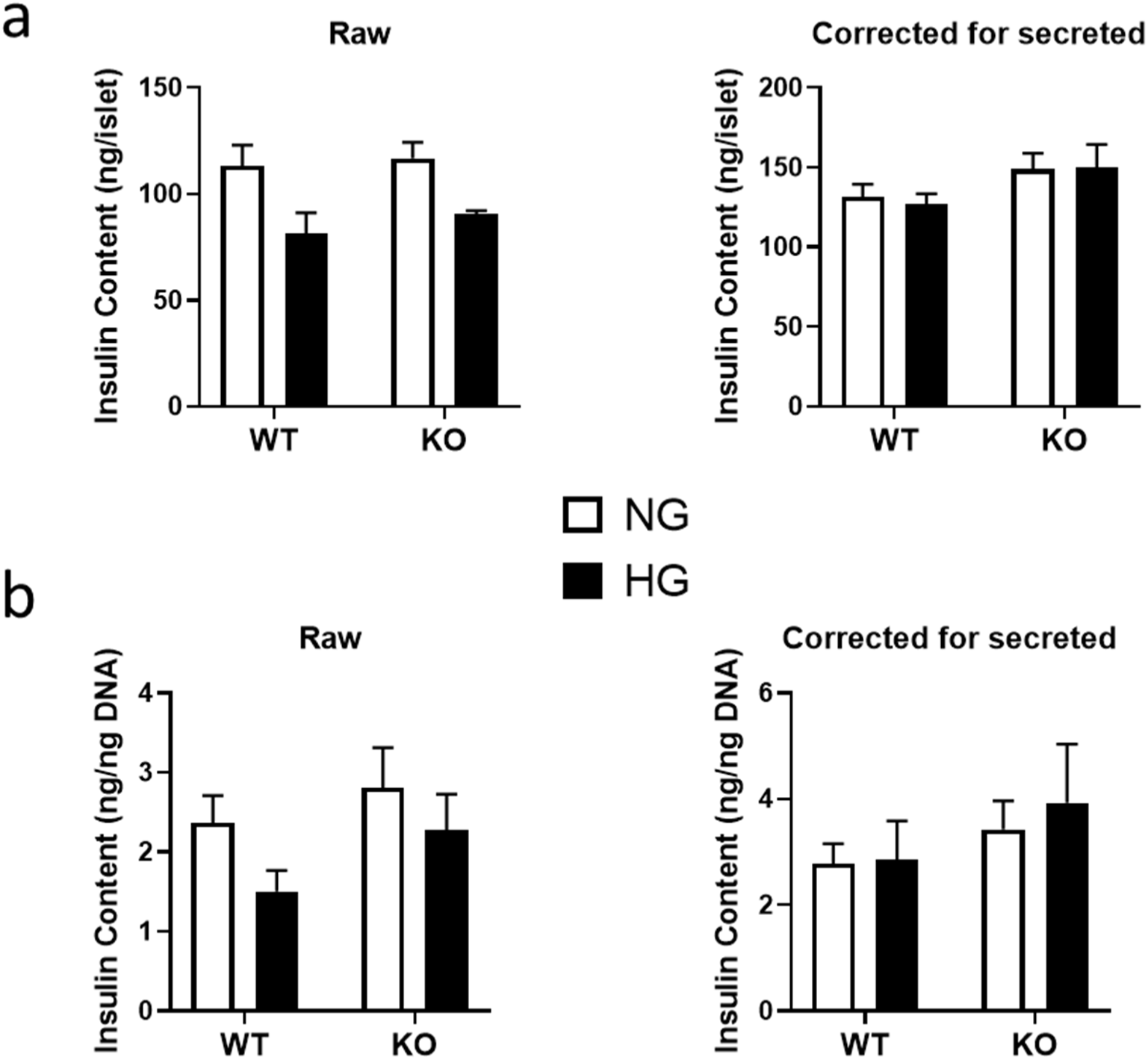
Insulin content in NG- and HG-cultured BclxβWT and BclxβKO islets. Insulin content was extracted and quantified from islets following ramp glucose-stimulated Insulin secretion assay (shown in Fig. 3). Extracted insulin content was expressed per islet (a) or per ng DNA (b) before and after correction for the total amount of insulin that was secreted during the glucose ramp. The tendency for HG-cultured islets to have lower insulin content is accounted for by their hyper-secretion of Insulin.

**Supplemental Figure 5 (related to Fig. 5).**
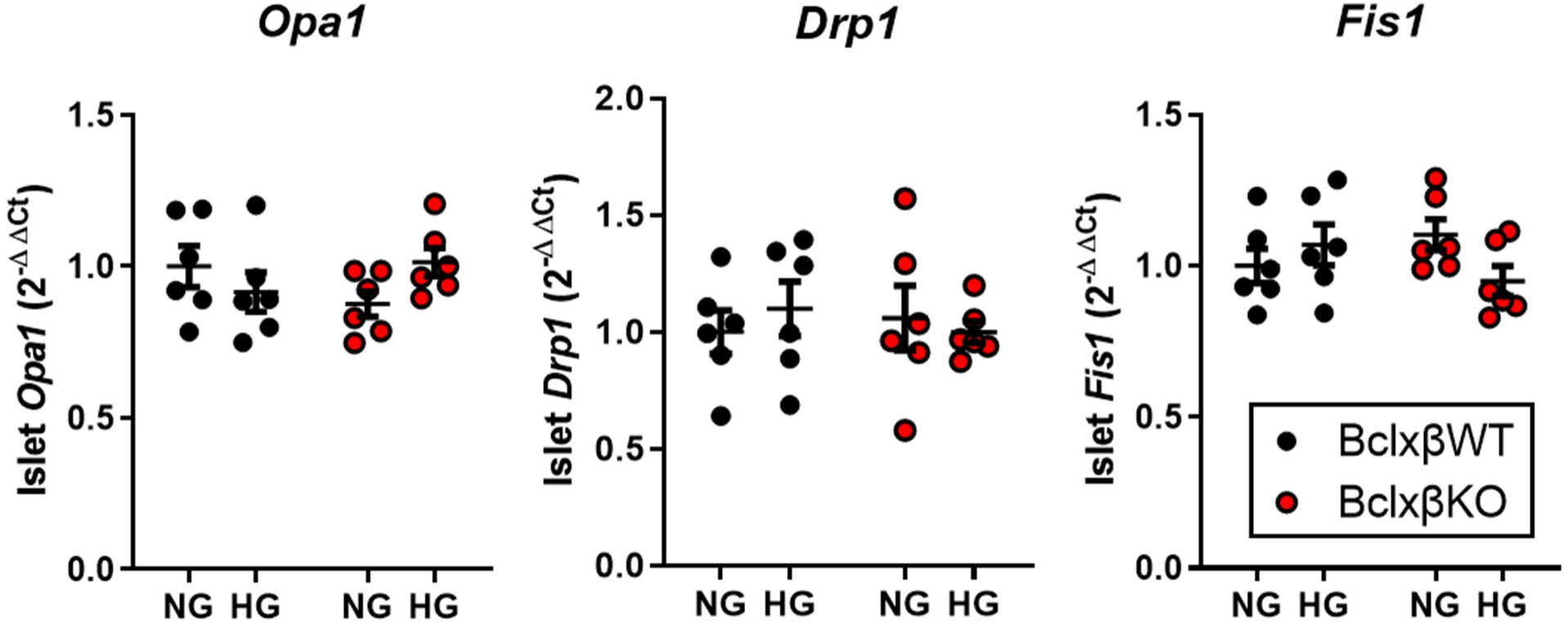
Expression of mitochondrial fusion & fission regulators. qPCR quantification of the expression of mitochondrial fusion (Opa1) and fission *(Drp1 & Fis1)* regulators in BclxβWT and BclxβKO islets after 6 days culture in NG or HG. No significant differences by two-way ANOVA followed by multiple comparisons tests (n=6).

**Supplemental Figure 6 (related to figure 7).**
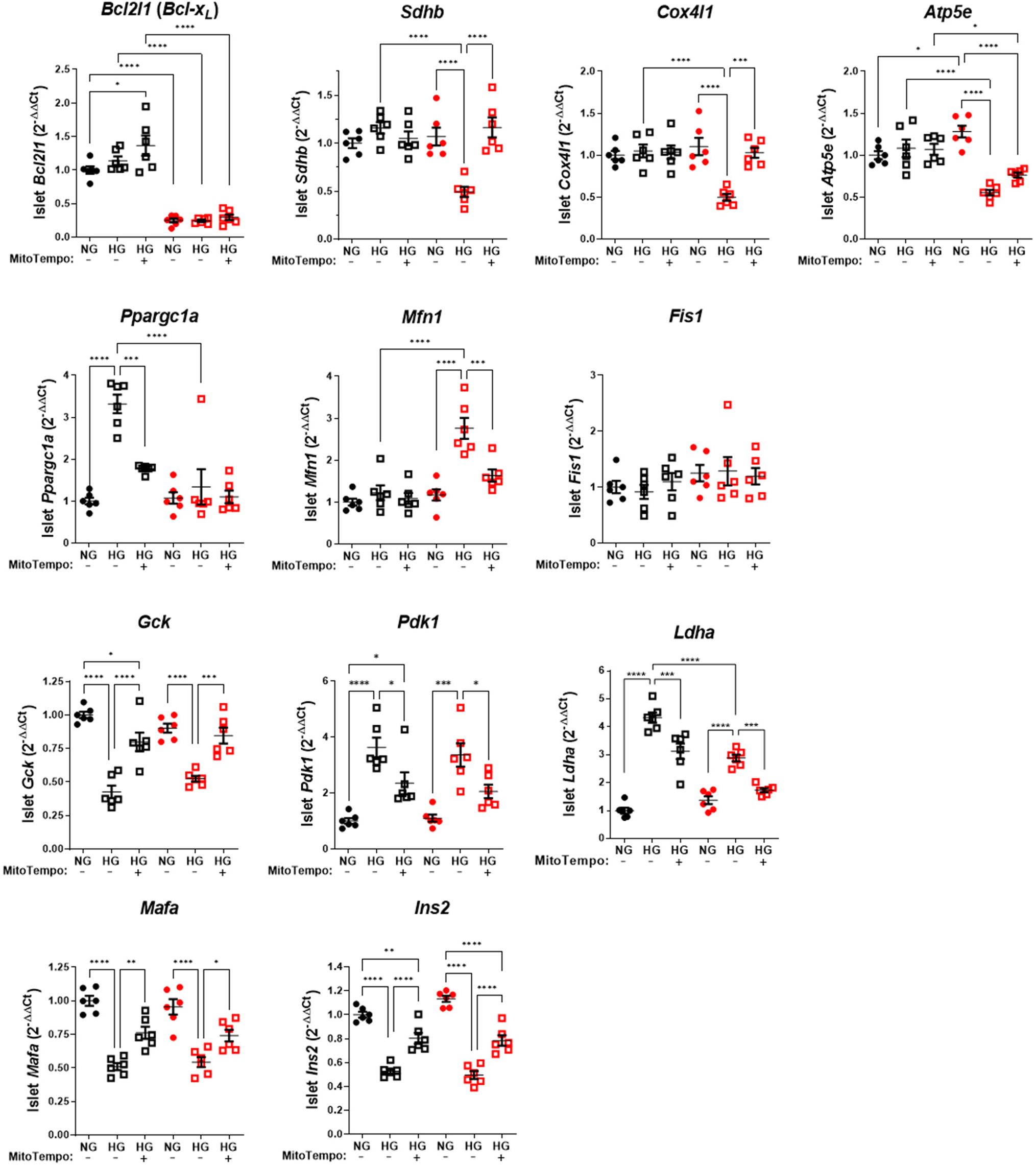
qPCR analyses showing that mitoROS drives loss of ETC gene expression, metabolic reprogramming and loss of identity in HG-cultured 13-cells. A heat map summary of these, and addit ional, qPCR experiments is shown in Fig. 7b.

## REFERENCES

1. Irles, E. et al. Enhanced glucose-induced intracellular signaling promotes insulin hypersecretion: pancreatic beta-cell functional adaptations in a model of genetic obesity and prediabetes. Mol Cell Endocrinol 404, 46–55 (2015).

2. Chen, C. et al. Alterations in beta-Cell Calcium Dynamics and Efficacy Outweigh Islet Mass Adaptation in Compensation of Insulin Resistance and Prediabetes Onset. Diabetes 65, 2676–85 (2016).

3. Prentki, M., Matschinsky, F.M. & Madiraju, S.R. Metabolic signaling in fuel-induced insulin secretion. Cell Metab 18, 162–85 (2013).

4. Anello, M. et al. Functional and morphological alterations of mitochondria in pancreatic beta cells from type 2 diabetic patients. Diabetologia 48, 282–9 (2005).

5. Dlaskova, A. et al. 4Pi microscopy reveals an impaired three-dimensional mitochondrial network of pancreatic islet beta-cells, an experimental model of type-2 diabetes. Biochim Biophys Acta 1797, 1327–41 (2010).

6. Haythorne, E. et al. Diabetes causes marked inhibition of mitochondrial metabolism in pancreatic beta-cells. Nat Commun 10, 2474 (2019).

7. Ebrahimi, A.G. et al. Beta cell identity changes with mild hyperglycemia: Implications for function, growth, and vulnerability. Mol Metab 35, 100959 (2020).

8. Hou, J. et al. Temporal Transcriptomic and Proteomic Landscapes of Deteriorating Pancreatic Islets in Type 2 Diabetic Rats. Diabetes (2017).

9. Chareyron, I. et al. Augmented mitochondrial energy metabolism is an early response to chronic glucose stress in human pancreatic beta cells. Diabetologia (2020).

10. Gross, A. & Katz, S.G. Non-apoptotic functions of BCL-2 family proteins. Cell Death Differ 24, 1348–1358 (2017).

11. Aharoni-Simon, M. et al. Bcl-2 Regulates Reactive Oxygen Species Signaling and a Redox-Sensitive Mitochondrial Proton Leak in Mouse Pancreatic beta-Cells. Endocrinology 157, 2270–81 (2016).

12. Luciani, D.S. et al. Bcl-2 and Bcl-xL Suppress Glucose Signaling in Pancreatic beta-Cells. Diabetes 62, 170–82 (2013).

13. Zhou, Y.P. et al. Overexpression of Bcl-x(L) in beta-cells prevents cell death but impairs mitochondrial signal for insulin secretion. Am J Physiol Endocrinol Metab 278, E340–51 (2000).

14. Danial, N.N. et al. Dual role of proapoptotic BAD in insulin secretion and beta cell survival. Nat Med 14, 144–53 (2008).

15. Ahn, S.H. et al. Tamoxifen suppresses pancreatic beta-cell proliferation in mice. PLoS One 14, e0214829 (2019).

16. Carboneau, B.A., Le, T.D., Dunn, J.C. & Gannon, M. Unexpected effects of the MIP-CreER transgene and tamoxifen on beta-cell growth in C57Bl6/J male mice. Physiol Rep 4(2016).

17. Luciani, D.S., Ao, P., Hu, X., Warnock, G.L. & Johnson, J.D. Voltage-gated Ca(2+) influx and insulin secretion in human and mouse beta-cells are impaired by the mitochondrial Na(+)/Ca(2+) exchange inhibitor CGP-37157. Eur J Pharmacol 576, 18–25 (2007).

18. Brand, M.D. & Nicholls, D.G. Assessing mitochondrial dysfunction in cells. Biochem J 435, 297–312 (2011).

19. Chaudhry, A., Shi, R. & Luciani, D.S. A pipeline for multidimensional confocal analysis of mitochondrial morphology, function, and dynamics in pancreatic beta-cells. Am J Physiol Endocrinol Metab 318, E87–E101 (2020).

20. Chaudhry, A. & Luciani, D.S. Mitochondria Analyzer. (https://github.com/AhsenChaudhry/Mitochondria-Analyzer, 2019).

21. Hemel, I., Engelen, B.P.H., Luber, N. & Gerards, M. A hitchhiker’s guide to mitochondrial quantification. Mitochondrion 59, 216–224 (2021).

22. Bolger, A.M., Lohse, M. & Usadel, B. Trimmomatic: a flexible trimmer for Illumina sequence data. Bioinformatics 30, 2114–20 (2014).

23. Patro, R., Duggal, G., Love, M.I., Irizarry, R.A. & Kingsford, C. Salmon provides fast and bias-aware quantification of transcript expression. Nat Methods 14, 417–419 (2017).

24. Soneson, C., Love, M.I. & Robinson, M.D. Differential analyses for RNA-seq: transcript-level estimates improve gene-level inferences. F1000Res 4, 1521 (2015).

25. Love, M.I., Huber, W. & Anders, S. Moderated estimation of fold change and dispersion for RNA-seq data with DESeq2. Genome Biol 15, 550 (2014).

26. Zhu, A., Ibrahim, J.G. & Love, M.I. Heavy-tailed prior distributions for sequence count data: removing the noise and preserving large differences. Bioinformatics 35, 2084–2092 (2019).

27. Huang da, W., Sherman, B.T. & Lempicki, R.A. Bioinformatics enrichment tools: paths toward the comprehensive functional analysis of large gene lists. Nucleic Acids Res 37, 1–13 (2009).

28. Huang da, W., Sherman, B.T. & Lempicki, R.A. Systematic and integrative analysis of large gene lists using DAVID bioinformatics resources. Nat Protoc 4, 44–57 (2009).

29. Luciani, D.S. et al. Roles of IP_3_R and RyR Ca^2+^ channels in endoplasmic reticulum stress and beta-cell death. Diabetes 58, 422–32 (2009).

30. Chan, J.Y. et al. The balance between adaptive and apoptotic unfolded protein responses regulates beta-cell death under ER stress conditions through XBP1, CHOP and JNK. Mol Cell Endocrinol 413, 189–201 (2015).

31. Federici, M. et al. High glucose causes apoptosis in cultured human pancreatic islets of Langerhans: a potential role for regulation of specific Bcl family genes toward an apoptotic cell death program. Diabetes 50, 1290–301 (2001).

32. Bensellam, M., Montgomery, M.K., Luzuriaga, J., Chan, J.Y. & Laybutt, D.R. Inhibitor of differentiation proteins protect against oxidative stress by regulating the antioxidant-mitochondrial response in mouse beta cells. Diabetologia 58, 758–70 (2015).

33. Tanaka, Y., Tran, P.O., Harmon, J. & Robertson, R.P. A role for glutathione peroxidase in protecting pancreatic beta cells against oxidative stress in a model of glucose toxicity. Proc Natl Acad Sci U S A 99, 12363–8 (2002).

34. Bensellam, M. et al. Phlda3 regulates beta cell survival during stress. Sci Rep 9, 12827 (2019).

35. Kale, J., Osterlund, E.J. & Andrews, D.W. BCL-2 family proteins: changing partners in the dance towards death. Cell Death Differ (2017).

36. Pullen, T.J. et al. Identification of genes selectively disallowed in the pancreatic islet. Islets 2, 89–95 (2010).

37. Pullen, T.J. & Rutter, G.A. When less is more: the forbidden fruits of gene repression in the adult beta-cell. Diabetes Obes Metab 15, 503–12 (2013).

38. Kim-Muller, J.Y. et al. Aldehyde dehydrogenase 1a3 defines a subset of failing pancreatic beta cells in diabetic mice. Nat Commun 7, 12631 (2016).

39. Kuo, T. et al. Induction of alpha cell-restricted Gc in dedifferentiating beta cells contributes to stress-induced beta-cell dysfunction. JCI Insight 5(2019).

40. Mookerjee, S.A., Gerencser, A.A., Nicholls, D.G. & Brand, M.D. Quantifying intracellular rates of glycolytic and oxidative ATP production and consumption using extracellular flux measurements. J Biol Chem 292, 7189–7207 (2017).

41. Kukat, C. et al. Cross-strand binding of TFAM to a single mtDNA molecule forms the mitochondrial nucleoid. Proc Natl Acad Sci U S A 112, 11288–93 (2015).

42. Gauthier, B.R. et al. PDX1 deficiency causes mitochondrial dysfunction and defective insulin secretion through TFAM suppression. Cell Metab 10, 110–8 (2009).

43. Silva, J.P. et al. Impaired insulin secretion and beta-cell loss in tissue-specific knockout mice with mitochondrial diabetes. Nat Genet 26, 336–40 (2000).

44. Eisner, V., Picard, M. & Hajnoczky, G. Mitochondrial dynamics in adaptive and maladaptive cellular stress responses. Nat Cell Biol 20, 755–765 (2018).

45. Chan, D.C. Mitochondrial Dynamics and Its Involvement in Disease. Annu Rev Pathol (2019).

46. Pearson, G.L., Gingerich, M.A., Walker, E.M., Biden, T.J. & Soleimanpour, S.A. A Selective Look at Autophagy in Pancreatic beta-Cells. Diabetes 70, 1229–1241 (2021).

47. Ploumi, C., Daskalaki, I. & Tavernarakis, N. Mitochondrial biogenesis and clearance: a balancing act. FEBS J 284, 183–195 (2017).

48. Berman, S.B. et al. Bcl-x L increases mitochondrial fission, fusion, and biomass in neurons. J Cell Biol 184, 707–19 (2009).

49. Campbell, J.E. & Newgard, C.B. Mechanisms controlling pancreatic islet cell function in insulin secretion. Nat Rev Mol Cell Biol 22, 142–158 (2021).

50. Willems, P.H., Rossignol, R., Dieteren, C.E., Murphy, M.P. & Koopman, W.J. Redox Homeostasis and Mitochondrial Dynamics. Cell Metab 22, 207–18 (2015).

51. Quiros, P.M., Mottis, A. & Auwerx, J. Mitonuclear communication in homeostasis and stress. Nat Rev Mol Cell Biol 17, 213–26 (2016).

52. da Cunha, F.M., Torelli, N.Q. & Kowaltowski, A.J. Mitochondrial Retrograde Signaling: Triggers, Pathways, and Outcomes. Oxid Med Cell Longev 2015, 482582 (2015).

53. Picard, M., Shirihai, O.S., Gentil, B.J. & Burelle, Y. Mitochondrial morphology transitions and functions: implications for retrograde signaling? Am J Physiol Regul Integr Comp Physiol 304, R393–406 (2013).

54. Laybutt, D.R. et al. Critical reduction in beta-cell mass results in two distinct outcomes over time. Adaptation with impaired glucose tolerance or decompensated diabetes. J Biol Chem 278, 2997–3005 (2003).

55. Jonas, J.C. et al. Chronic hyperglycemia triggers loss of pancreatic beta cell differentiation in an animal model of diabetes. J Biol Chem 274, 14112–21 (1999).

56. White, S.A., Zhang, L.S., Pasula, D.J., Yang, Y.H.C. & Luciani, D.S. Bax and Bak jointly control survival and dampen the early unfolded protein response in pancreatic beta-cells under glucolipotoxic stress. Sci Rep 10, 10986 (2020).

57. Hetz, C. et al. Proapoptotic BAX and BAK modulate the unfolded protein response by a direct interaction with IRE1alpha. Science 312, 572–6 (2006).

58. Molina, A.J. et al. Mitochondrial networking protects beta-cells from nutrient-induced apoptosis. Diabetes 58, 2303–15 (2009).

59. Oropeza, D. et al. PGC-1 coactivators in beta-cells regulate lipid metabolism and are essential for insulin secretion coupled to fatty acids. Mol Metab 4, 811–22 (2015).

60. Twig, G. et al. Fission and selective fusion govern mitochondrial segregation and elimination by autophagy. EMBO J 27, 433–46 (2008).

61. Alavian, K.N. et al. Bcl-x(L) regulates metabolic efficiency of neurons through interaction with the mitochondrial F(1)F(O) ATP synthase. Nat Cell Biol (2011).

